# Respiration and brain neural dynamics associated with interval timing during odor fear learning in rats

**DOI:** 10.1101/2020.07.31.228866

**Authors:** Maryne Dupin, Samuel Garcia, Belkacem Messaoudi, Valérie Doyère, Anne-Marie Mouly

## Abstract

In fear conditioning, where a conditioned stimulus predicts the arrival of an aversive stimulus, the animal encodes the time interval between the two stimuli. Freezing, the most used index to assess learned fear, lacks the temporal resolution required to investigate interval timing at the early stages of learning. Here we monitored respiration to visualize anticipatory behavioral responses in an odor fear conditioning in rats, while recording theta (5-15Hz) and gamma (40-80Hz) brain oscillatory activities in the medial prefrontal cortex (mPFC), basolateral amygdala (BLA), dorsomedial striatum (DMS) and olfactory piriform cortex (PIR). We investigated the temporal patterns of respiration frequency and of theta and gamma activity power during the odor-shock interval. We found that akin to respiration patterns, theta temporal curves were modulated by the duration of the odor-shock interval in the four recording sites, and respected scalar property in mPFC and DMS. In contrast, gamma temporal curves were modulated by the interval duration only in the mPFC, and in a manner that did not respect scalar property. This suggests a preferential role for theta rhythm in interval timing. In addition, our data bring the novel idea that the respiratory rhythm might take part in the setting of theta activity dynamics.

## INTRODUCTION

Interval timing refers to the ability to time intervals ranging from seconds to minutes and guides fundamental animal behaviors like the anticipation of rewarding or aversive events. The tasks classically used in the literature to assess interval timing in animals necessitate numerous training sessions and involve a motor response from the animal (Roberts, 1981). Yet some studies show that in associative learning, animals learn to time the arrival of reinforcement from the outset of conditioning (Davis et al., 1989; Arcediano et al., 2003; Drew et al., 2005; Diaz-Mataix et al., 2013; Shionoya et al, 2013; Boulanger-Bertolus et al, 2014) and such temporal encoding has been suggested to be a fundamental component of associative learning (Balsam and Gallistel, 2009). However, the neurobiological basis of interval timing in Pavlovian associative learning remain poorly understood, due in part to the paucity of studies designed for its investigation (Tallot and Doyère, 2020).

In a previous study using odor fear conditioning in rats, we showed that, when using an appropriate index, namely the respiratory rate, interval timing can be inferred from the animal’s behavior after a few training trials (Shionoya et al, 2013). More specifically, the animal’s respiratory rate was monitored in this paradigm where an initially neutral odor signals the arrival of an aversive mild foot-shock at a fixed time interval. We showed that after a few odor-shock pairings, the respiratory frequency curve presented a temporal pattern that was linked to the duration of the interval to be timed, in a manner that respected scalar property, a hallmark of interval timing, i.e. the error magnitude in estimating a duration was proportional to the duration to be timed (Gibbon and Church, 1990).

Based on these findings, in the present study, we investigated the neural network dynamics occurring during the odor-shock interval in odor fear conditioning in rats. Although the neural mechanisms underlying timing remain largely unknown, several studies in the literature reported dynamically changing patterns of activity that contain information about elapsed time since a given stimulus. This property has been found in multiple brain areas including the dorsal striatum and the prefrontal cortex (Coull et al, 2011; Buhusi and Meck, 2005). Recent studies using aversive conditioning suggest that the amygdala also plays a role in interval timing (Diaz-Mataix et al, 2013; Shionoya et al, 2013; Dallérac et al, 2017; Tallot et al, 2020) and regulates physiological correlates of plasticity in the striatum, thanks to direct amygdala projections to the striatum (Dallérac et al, 2017). In addition, evidence of time encoding has been found in different sensory cortices like the primary visual cortex (Shuler and Bear, 2006; Bueti et al, 2010) and the olfactory piriform cortex (Hegoburu et al, 2009). Therefore, in the present study, rats were implanted with chronic electrodes in four brain areas: the median prefrontal cortex (mPFC), basolateral amygdala (BLA), dorsomedial striatum (DMS) and piriform cortex (PIR). The animals were trained in an odor fear conditioning paradigm while their respiration was measured, together with brain oscillatory activities in the targeted network. We focused on variations in the theta (5-15Hz) and gamma (40-80Hz) bands that have been shown to reflect temporal processing (Headley and Weinberger, 2011; Dallérac et al, 2017; Popescu et al, 2009; Parker et al, 2014).

Our prediction was that, if a brain area is involved in timing the odor-shock interval, temporal curves of oscillatory activity power during that interval should 1) build up after a few odor-shock pairings in parallel to respiratory frequency temporal curves, and 2) be modulated by the duration of the interval in a manner that respects scalar property.

## RESULTS

### Respiratory frequency is a good index of interval timing providing the level of stress is not too high

In a previous study (Shionoya et al, 2013) we showed that respiration is a good index of interval timing in non-implanted animals. We assessed if this was still true for implanted animals, in which the level of stress could be higher and might constrain the range of the respiratory response.

The time course of respiratory frequency during the odor-shock interval in Group 20s was compared between the Conditioning session where a 20s interval was used (see protocol in Figure 1), and the Shift session for which the interval was shifted to 30s (Figure 2A). The ANOVA revealed no significant effect of Interval, but a significant effect of Time (F_17,340_=11.43, p<0.0001) and a significant Interval×Time interaction (F_17,340_=3.029, p<0.0001). As depicted on Figure 2A, shifting from a 20s interval to a 30s interval resulted in an increase of the mean latency to the peak respiratory frequency on the temporal curve (5s for 20s interval vs 8s for 30s interval).

**Figure 1:**
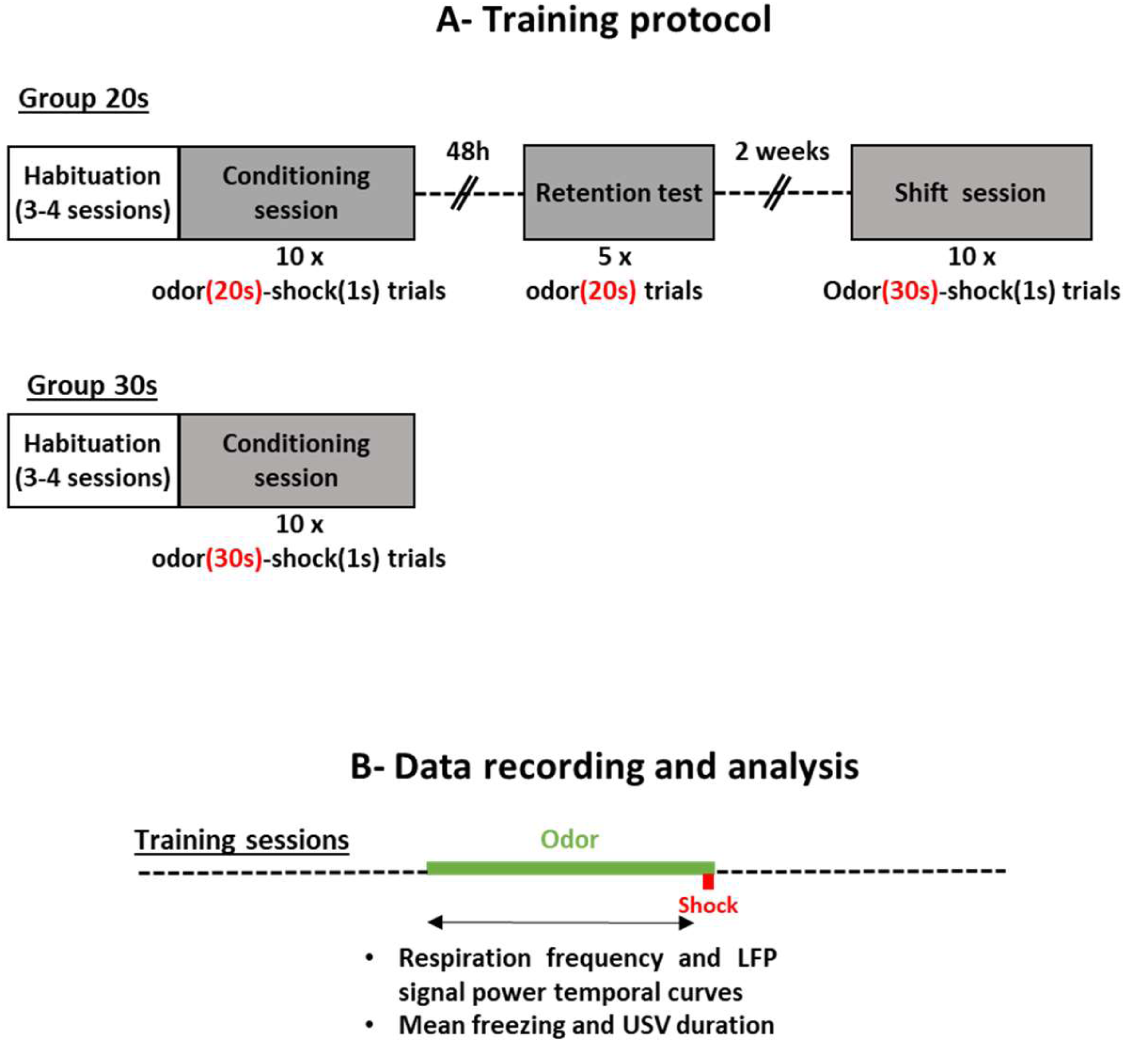
A) Training protocol. Group 20s: The animals were trained with 10 Odor-Shock pairings using a 20s Odor-Shock interval (Conditioning session). 48h hours later, they were tested for their retention of the learning (Retention test) with 5 presentations of the learned odor. Two weeks later they were trained with 10 Odor-Shock pairings using a 30s Odor-Shock interval (Shift session). Group 30s: The animals were trained with 10 Odor-Shock pairings using a 30s Odor-Shock interval (Conditioning session). B) Data recording and analysis. Training sessions: in the two groups, during the odor-shock interval, local field potentials (LFP), respiration, ultrasonic vocalizations and freezing behavior were recorded, synchronized and analyzed as described in the Methods.

**Figure 2:**
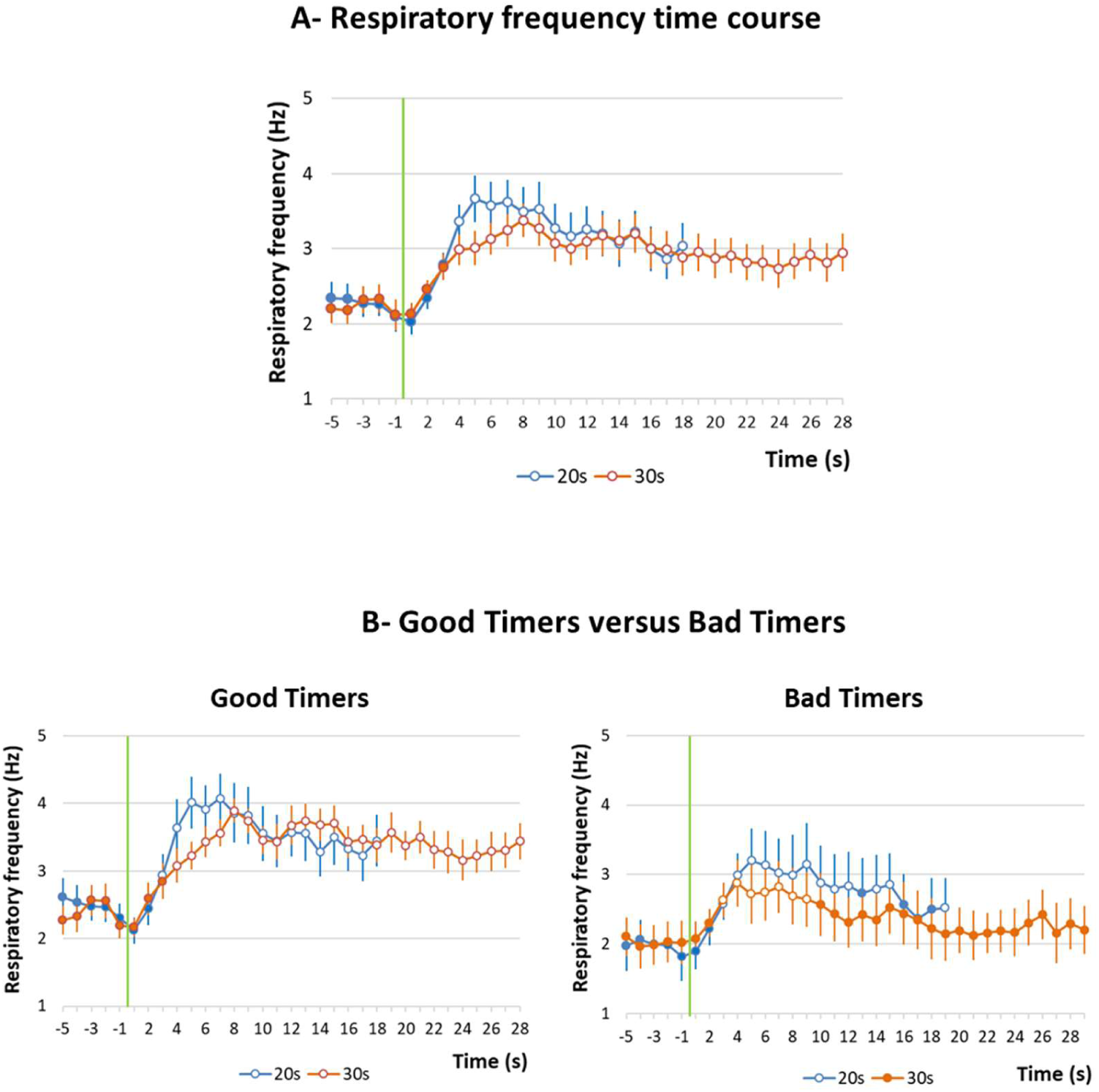
A) Respiratory frequency time course in Group 20s. The time course of respiratory frequency is represented with a 1-s bin precision, from 5s before odor onset (green vertical line) to shock arrival which occurred either at 20s (Conditioning session, in blue) or at 30s (Shift session 2, in orange). B) Good Timers versus Bad Timers subgroups. In Good Timers subgroup, the temporal patterns of respiratory frequency were modulated by the duration of the odor-shock interval, whereas in Bad Timers subgroup, they were not. Open circles on each curve signal the values that are significantly different from the baseline (p≤0.05).

However, based on individual temporal curves inspection, it occurred that two subgroups of animals could be distinguished: those presenting clearly shifted temporal curves for 20s and 30s intervals, and those presenting similar curves for both intervals. To quantify it, we measured for each animal the time at which the respiratory frequency peaked for the 20s- and 30s-interval temporal curves and calculated the difference between the two obtained values. When this value was ≥2sec, the animal was classified as a Good Timer, the remaining animals were classified as Bad Timers. This led to two subgroups: subgroup Good Timers (n=12, mean latency difference = 6.08±0.96 sec) and subgroup Bad Timers (n=9, mean latency difference = −2.22±1.00 sec). The temporal curves obtained in these two subgroups are presented in Figure 2B. In the Good Timers subgroup (Figure 2B, left), the ANOVA confirmed a significant effect of Time (F_17,187_=16.31, p<0.0001) and a significant Interval×Time interaction (F_17,187_=3.09, p<0.0001), with no significant effect of Interval. A significant interaction between interval duration and elapsed time indicates that the time course of the respiratory frequency changes is affected by the duration of the odor-shock interval. In the Bad Timers subgroup (Figure 2B, right), the ANOVA revealed a significant effect of Time (F_17,136_=3.59, p<0.0001) but no significant effect of Interval and no significant Interval×Time interaction. Interestingly, the difference between Good Timers and Bad Timers is mainly due to the temporal patterns exhibited during the Shift session. Indeed the temporal patterns displayed by both subgroups during the Conditioning session (20s interval) were similar (GroupxTime interaction: F_17,323_ = 0.94, p=0.53), while those displayed during the Shift session (30s interval) differed significantly (GroupxTime interaction: F_27,513_ = 3.07, p<0.001).

The temporal patterns observed in Good Timers subgroup suggest that the Odor-Shock interval was timed according to the scalar rule (Gibbon, 1977). In order to assess scalar timing quantitatively, the time axis for the 30-s data was multiplicatively rescaled so that the 18 time-bins during the interval for both groups represented the same proportions of elapsed time from odor onset to shock presentation. The scalar timing rule predicts superior superposition of the functions in relative time, compared to no rescaling. Figure S1 shows raw and rescaled mean curves for both 20- and 30-s conditions, for Good Timers and Bad Timers subgroups. Superposition was indexed by eta-squared (η^2^), a measure of the proportion of variance accounted for by the mean of the two functions (Brown et al., 1992). When superposition is perfect, η^2^ is at its maximum value of 1. For Good Timers, η^2^ was greater under the multiplicative transform (η^2^=0.93, Supplementary Fig. S1-A, right) than under no rescaling (η^2^=0.88, Supplementary Fig. S1-A, left). In contrast, for Bad Timers subgroup, η2 was greater under no rescaling (η^2^=0.73, Supplementary Fig. S1-B, left) than under the multiplicative transform (η^2^=0.50, Supplementary Fig. S1-B, right).

We next evaluated to what extent the timing abilities assessed through respiration could be related to different levels of fear exhibited by the animals. To do so, we compared the mean amount of freezing exhibited by the animals in the Good Timers and Bad Timers subgroups during the Odor-Shock interval in the Conditioning and Shift sessions (Figure 3A, Left). The ANOVA revealed no significant effect of Subgroup, Training session, and Training session×Subgroup interaction. We then compared the number of USV emitted by the animals in the two subgroups during the Odor-Shock interval for the two training sessions. The ANOVA revealed a significant effect of Subgroup (F_1,16_=8.59, p=0.010) and Training session (F_1,16_=7.91, p=0.013), and no significant Training session×Subgroup interaction. Further pairwise comparisons showed that Bad Timers animals emitted more USV than Good Timers during both training sessions (p=0.012 for the 20s interval session; p=0.015 for the 30s interval session).

**Figure 3:**
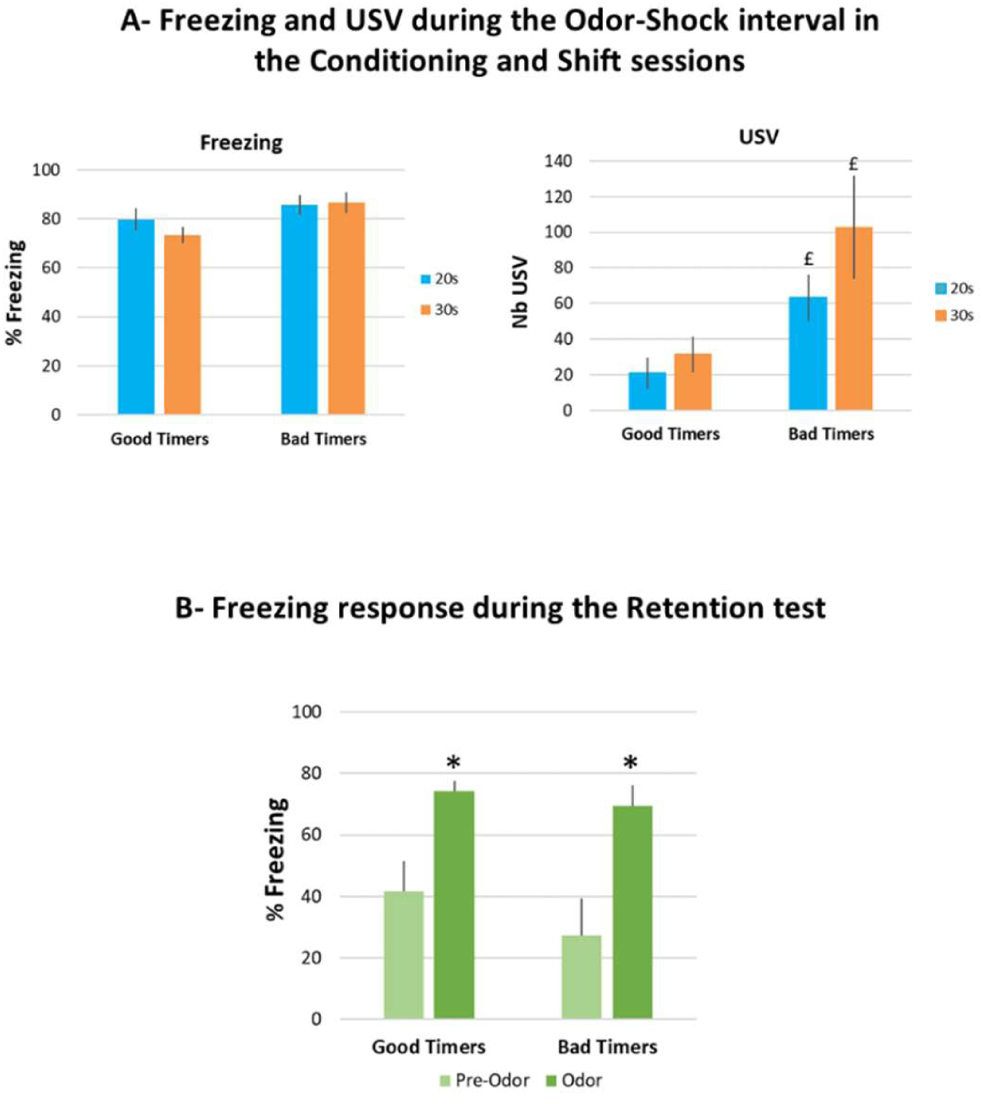
A) Freezing and ultrasonic vocalizations (USV) during the two training sessions, in Good Timers and Bad Timers subgroups. Freezing amount and number of USVs were measured during the 20s (Conditioning session, in blue) and 30s (Shift session, in orange) odor-shock intervals throughout the training session. £: significant between group difference (p≤0.05). B) Freezing amount during the Retention test. Good Timers and Bad Timers subgroups showed similar levels of freezing during the Retention test. *: significant within group difference with pre-odor period (p≤0.05).

Finally, we assessed whether the two subgroups presented different levels of learned fear during the 48h retention test (Figure 3B). In each subgroup, we compared the amounts of freezing observed before odor introduction and during odor. The ANOVA revealed a significant effect of Period (Pre-Odor vs Odor; F_1,18_=35.56, p<0.0001) but no significant effect for Subgroup nor for PeriodxSubgroup interaction. In both subgroups, the amount of freezing was higher in presence of the learned odor than before its delivery suggesting similar levels of fear memory in Bad Timers and Good Timers animals. Together, these data show that the temporal pattern observed in respiration rate in the Good Timers subgroup reflects the learning of Odor-Shock interval duration, and confirm our previous finding that rats can learn the expected time of shock delivery in an odor fear conditioning task within 10 conditioning trials (Shionoya et al, 2013). The ability to express temporal behavior related to interval duration, at least when looking at the respiratory rate, seems to depend on the animal’s fear response: the higher the USV amount, the lower the possibility for respiration to exhibit different temporal patterns related to interval duration. Finally, this difference in timing abilities seems to have no impact on the strength of the fear memory as assessed through odor-elicited freezing 48h later.

In the next analyses, we assessed whether the differences observed in timing expression in Bad Timers and Good Timers, based on their temporal pattern of respiratory frequency, were associated with similar differences in oscillatory activity temporal patterns.

### Changes in oscillatory activity in the theta (5-15Hz) band

We assessed whether, akin to respiratory frequency, the temporal patterns of theta activity power was modulated by the duration of the Odor-Shock interval.

As explained in the Methods, after elimination of artifacted electrophysiological signals, the number of animals per recording site were the following: BLA, n=17; CPF, n=20; PIR, n=20; DMS, n=19 for Group 20s; BLA, n=6; CPF, n=8; PIR, n=8; DMS, n=9 for Group 30s.

The temporal patterns of theta activity power during the Odor-Shock interval in Group 20s was compared between the Conditioning session where a 20s interval was used, and the Shift session for which the interval was shifted to 30s. Figure 4 presents the temporal curves obtained in each of the four recording sites for the Good Timers (left side) and Bad Timers (right side) subgroups. The ANOVA carried out in the Good Timers subgroup revealed no significant effect of Interval, but for all four structures a significant effect of Time (mPFC: F_17,187_=6.56, p<0.0001; BLA: F_17,119_=6.77, p<0.0001; PIR: F_17,187_=4.15, p<0.0001; DMS: F_17,187_=6.24, p<0.0001), and a significant IntervalxTime interaction (mPFC: F_17,187_=2.66, p<0.0001; BLA: F_17,119_=5.14, p<0.0001; PIR: F_17,187_=3.47, p<0.0001; DMS: F_17,187_=4.26, p<0.0001). In contrast, in the Bad Timers subgroup (Figure 4, right panels), theta activity power showed almost no change during the interval compared to baseline in the four recording sites (no significant effect of Time or Interval), and only the mPFC showed a significant Interval×Time interaction (F_17,119_=1.72, p=0.047).

**Figure 4:**
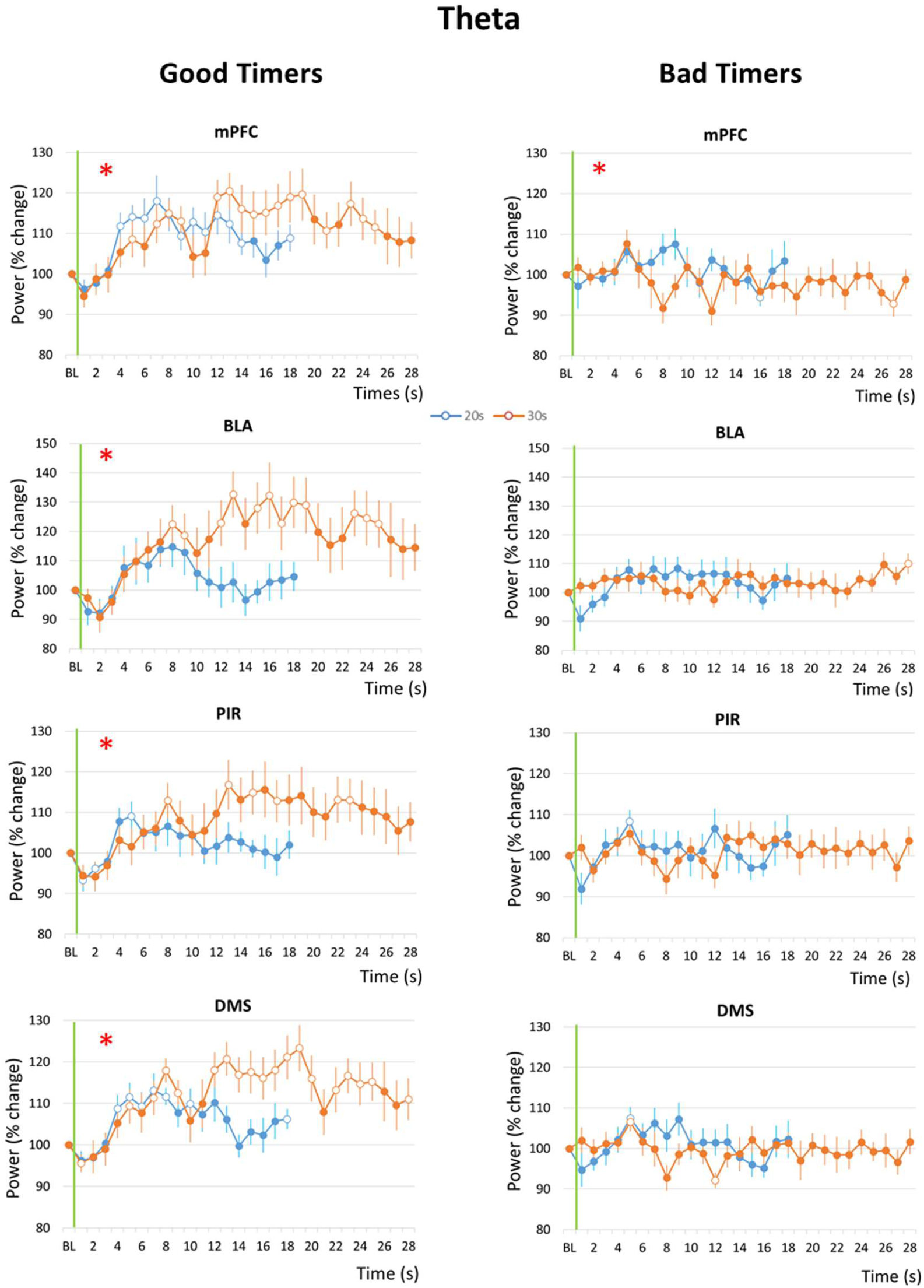
Theta (5-15Hz) band mean power variations during the odor-shock interval in the Good Timers (left side) and Bad Timers (right side) subgroups in the four recording sites. mPFC: medial prefrontal cortex; BLA: basolateral amygdala; PIR: olfactory piriform cortex; DMS: dorso-medial striatum. For each graph, theta power was expressed as percent change from the baseline (BL), and represented from odor onset (green vertical line) to shock arrival which occurs either at 20s (Conditioning session, in blue) or at 30s (Shift session, in orange). The red asterisk in the upper left corner of a graph signals a significant time x interval duration interaction. Open circles on each curve signal the values that are significantly different from the baseline (p≤0.05).

These data suggest that in Good Timers, the time course of theta activity power was modulated by the duration of the interval to be timed. In order to assess scalar timing quantitatively, the time axis for the 30-s data was multiplicatively rescaled and the η^2^ was calculated for each recording site (Supplementary Fig. S2). In mPFC and DMS, η^2^ was greater under the multiplicative transform (Supplementary Fig. S2, right) than under no rescaling (Supplementary Fig. S2, left), while the reverse was observed for BLA and PIR.

These data show that in Good Timers animals, akin to what is observed for respiration frequency, the time course of theta activity power was related to the learned duration of Odor-Shock interval in the four recording sites, and in a manner that respected scalar property in the mPFC and the DMS.

### Changes in oscillatory activity in the gamma (40-80Hz) band

Similar comparisons were made for the temporal pattern of gamma activity during the Odor-Shock interval. Figure 5 presents the temporal curves obtained in the four recording sites in the Good Timers (Left side) and Bad Timers (right side) subgroups. The ANOVA carried out in the Good Timers subgroup revealed no significant effect of Interval, but a significant effect of Time in mPFC (F_17,187_=4.73, p<0.0001), PIR (F_17,187_=2.57, p=0.002) and DMS (F_17,187_=3.65, p<0.0001), and a significant IntervalxTime interaction only in mPFC (F_17,187_=2.51, p=0.001). In the Bad Timers subgroup, the ANOVA revealed a significant effect of Time in mPFC (F_17,119_=2.24, p=0.006), PIR (F_17,119_=1.95, p=0.02) and DMS (F_17,102_=2.22, p=0.007), but no significant effect of Interval nor Interval×Time interaction in the four recording sites.

**Figure 5:**
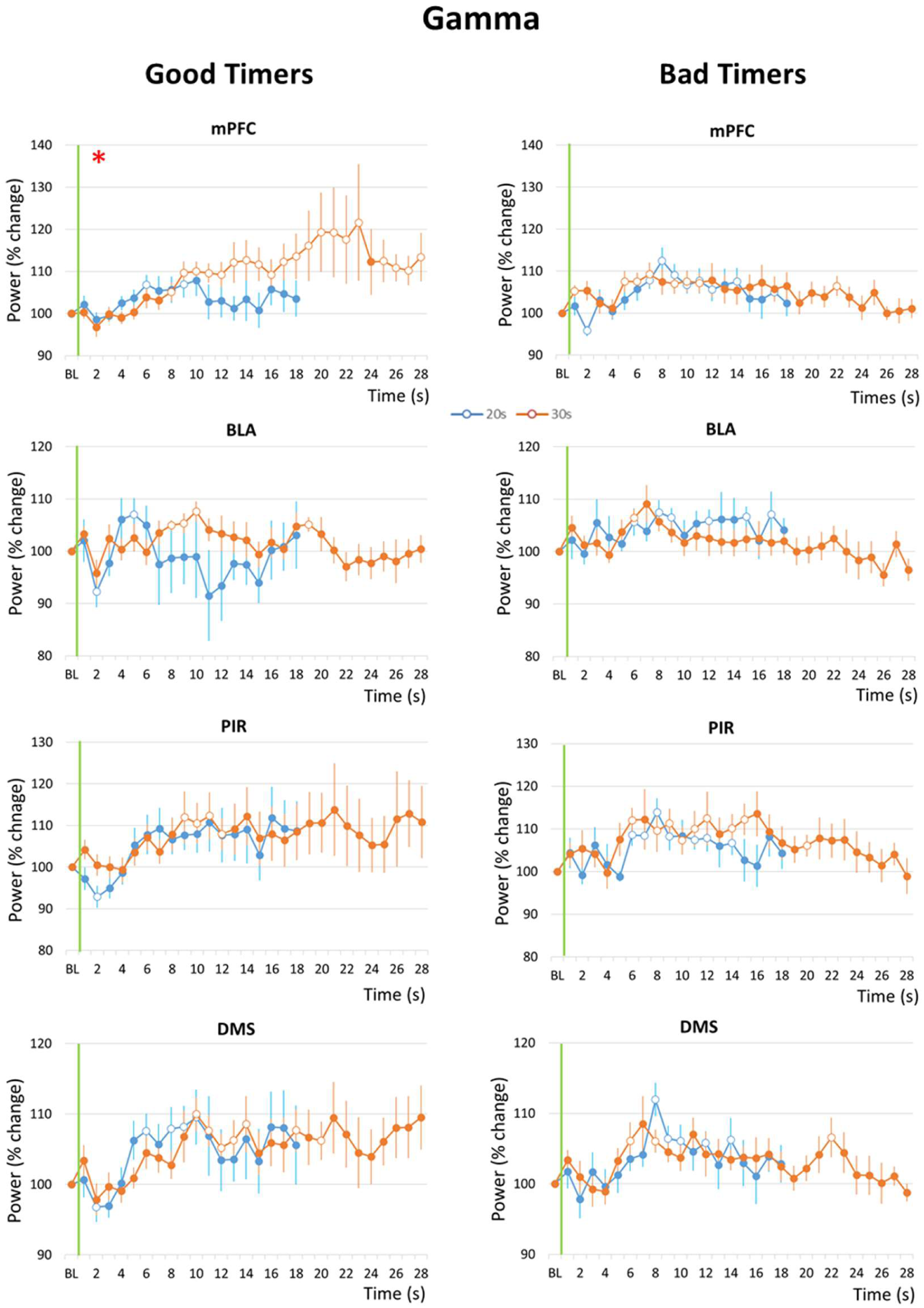
Gamma (40-80Hz) band mean power variations during the odor-shock interval in the Good Timers (left side) and Bad Timers (right side) subgroups in the four recording sites. mPFC: medial prefrontal cortex; BLA: basolateral amygdala; PIR: olfactory piriform cortex; DMS: dorso-medial striatum. For each graph, theta power was expressed as percent change from the baseline (BL), from the odor onset (green vertical line) to shock arrival which occurs either at 20s (Conditioning session, in blue) or at 30s (Shift session, in orange). The red asterisk in the upper left corner of a graph signals a significant time x interval duration interaction. Open circles on each curve signal the values that are significantly different from the baseline (p≤0.05).

Therefore, in both Good Timers and Bad Timers, gamma power increased significantly during the interval compared to baseline in the mPFC, DMS and PIR, but not in the BLA. When looking at the temporal patterns, the only structure in which the time course of gamma activity power was affected by the duration of the interval (i.e. showing an Interval x Time interaction) was the mPFC in the Good Timers subgroup. In order to assess scalar timing, the time axis for the 30-s data was multiplicatively rescaled and the η^2^ was calculated (Supplementary Fig. S3). η^2^ was greater under no rescaling (Supplementary Fig. S3, left) than under the multiplicative transform (Supplementary Fig. S3, right), suggesting gamma activity temporal pattern in mPFC does not solely reflect the learning of Odor-Shock interval duration.

### Correlation between respiration temporal curves and theta/gamma power temporal curves

Together the data presented so far suggest that in Good Timers, there might be a correlation between the time course of respiration frequency curve and that of theta activity power curves. To assess this quantitatively, we performed two kinds of correlation analysis. First, we measured for each animal the time of the peak respiratory frequency and the time of the peak theta activity power during the Conditioning session and calculated the correlation index between these values. This analysis showed significant positive correlation for BLA (R_8_ = 0.85, p<0.01), PIR (R_12_ = 0.63, p<0.02) and DMS (R_12_ = 0.61, p=0.02), and a tendency for mPFC (R_12_ = 0.53). We performed the same analysis for gamma activity and found no significant correlation with respiration (mPFC: R_12_ = 0.26, ns; BLA: R_8_ = 0.21, ns; PIR: R_12_ = 0.24, ns; DMS: R_12_ = −0.09, ns).

For the second correlation analysis, we calculated the difference between the times of the peak respiratory frequency during the Conditioning (20s interval) and Shift (30s interval) sessions: Peak 30s - Peak 20s (Shift R, Figure 6A). Similarly, for each recording site, we calculated the difference between the times of the peak theta activity power during the Conditioning and Shift sessions: Peak 30s - Peak 20s (Shift T, Figure 6B, upper part). Then we calculated the correlation between Shift R and Shift T (Figure 6B, lower part). The data show a significant positive correlation between the two values for the mPFC (R_12_ = 0.62, p<0.02), but not for the other sites (BLA: R_8_ = 0.43, ns; PIR: R_12_ =-0.49, ns; DMS: R_12_ = 0.32, ns). The same analysis was carried out for gamma activity in mPFC and showed no significant correlation (Shift G, Figure 6C, R_12_ = 0.39, ns).

**Figure 6:**
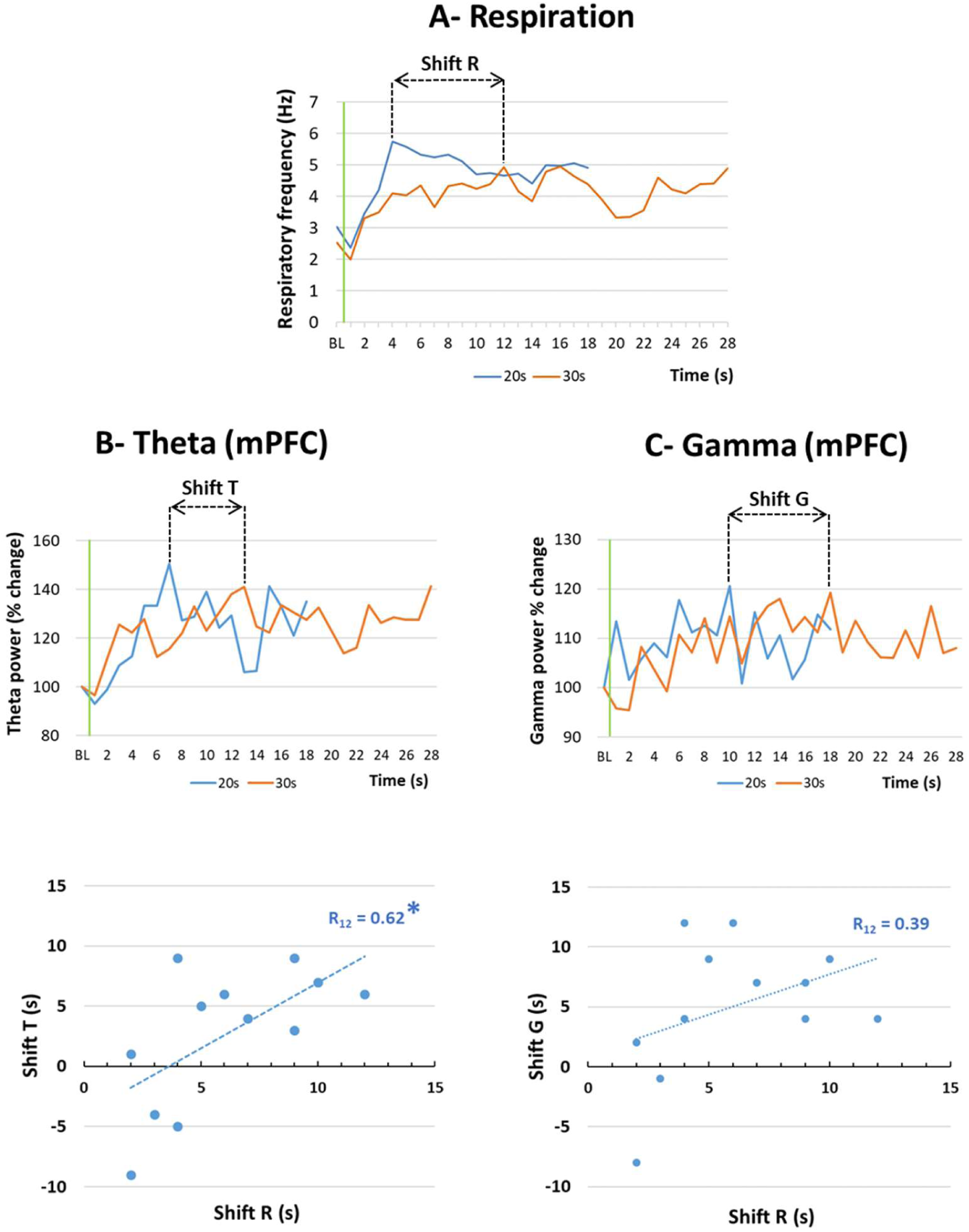
Correlation between the shift in respiration temporal curves and theta/gamma power temporal curves. A) The time of the peak respiratory frequency during the Conditioning session (20s interval) and the Shift session (30s intervals) was measured and the difference between the two values was used as the Shift R index. B) Upper graph: For each recording site, the time of the peak theta activity power during the Conditioning session (20s interval) and the Shift session (30s intervals) was measured and the difference between the two values was used as the Shift T index. The graph illustrates data from the mPFC: medial prefrontal cortex. Lower graph: the correlation between Shift R and Shift T indices was calculated. *significant correlation index, p≤0.05. C) Upper graph: For each recording site (example of the mPFC), the time of the peak gamma activity power during the Conditioning session (20s interval) and the Shift session (30s intervals) was measured and the difference between the two values was used as the Shift G index. Lower graph: the correlation between Shift R and Shift G indices was calculated.

In summary, in the four recorded structures, the time course of theta activity power correlates with the time course of respiratory frequency. In addition, in mPFC, the shift in theta activity power temporal curves observed when the interval duration was changed, was positively correlated with the shift in respiratory frequency temporal curve. These correlations were not observed when considering gamma activity power.

### Interval timing in Independent groups

In the experiments carried out so far, the same animals received two successive training sessions, a Conditioning session with a 20s odor-shock interval, and a Shift session with a 30s interval. This allowed us to compare the temporal curves within the same rats, thus discounting inter-individual variability. However, the data obtained during the Shift session might have been partly confounded by the previously experienced Conditioning session. In order to assess this point, a small group of animals was trained in a single session using a 30s odor-shock interval (Group 30s). We compared the temporal curves obtained in Good Timers during the Conditioning session of Group 20s to those of the single session of Group 30s. The only difference between the two groups was the duration of the odor-shock interval with an equivalent level of training.

Figure 7A (left part) presents the temporal curves obtained for respiratory frequency in the two groups. The ANOVA revealed no significant effect of Group, but a significant effect of Time (F_17,238_=5.39, p<0.001) and a significant Time×Group interaction (F_17,238_=2.01, p=0.012). When the time axis for the 30s data was multiplicatively rescaled to assess scalar timing (Figure 7A, right part), the η^2^ index increased from 0.56 (no rescaling) to 0.62 (multiplicative transform).

**Figure 7:**
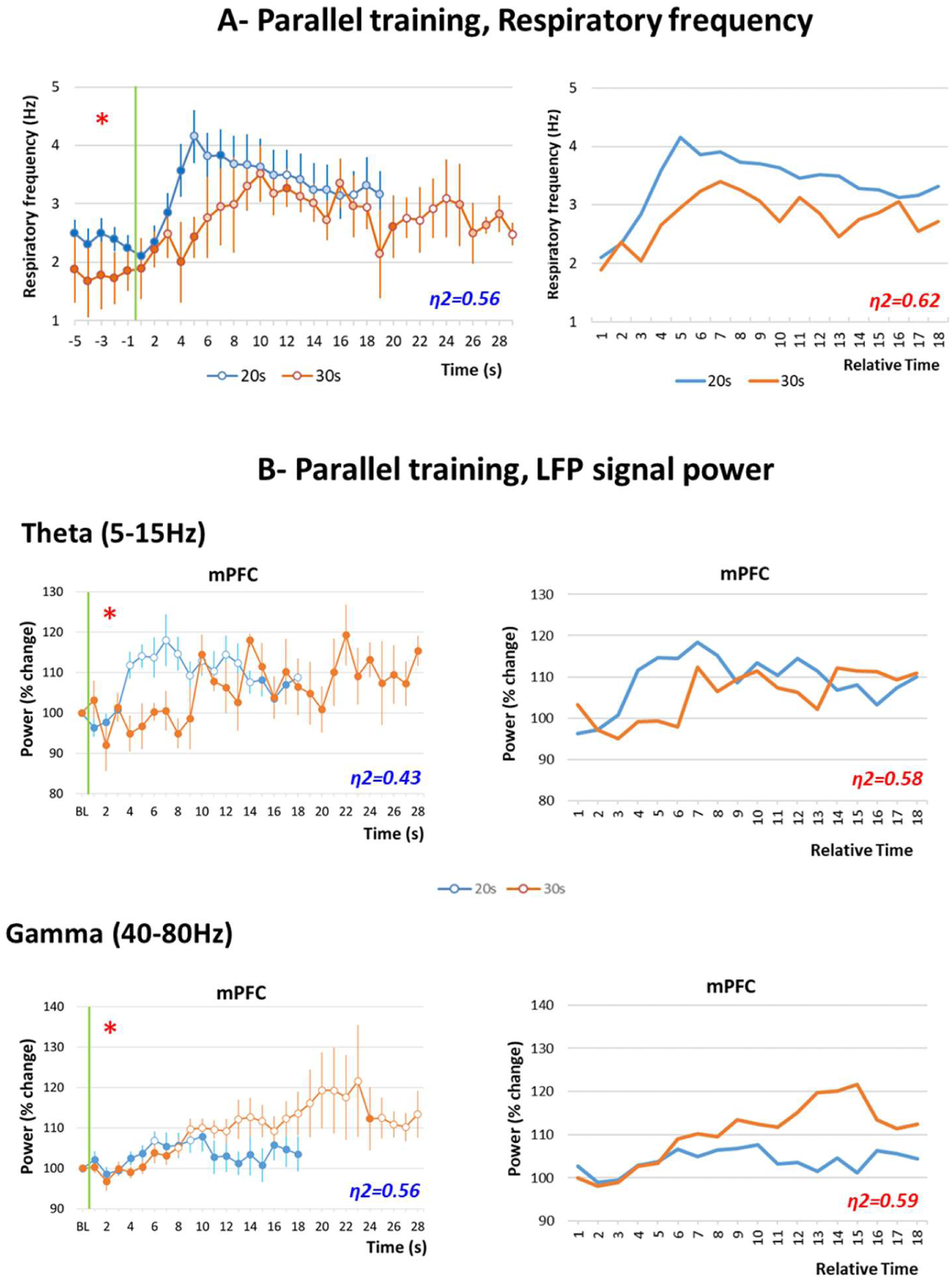
Behavioral and electrophysiological comparison in parallel training condition. A) Respiratory frequency curves. Left side, raw data: the temporal pattern of respiratory frequency is represented with a 1-s bin precision, from 5s before odor onset (green vertical line) to shock arrival which occurs either at 20s (Group 20s, in blue) or at 30s (Group 30s, in orange). Right side, rescaled curves: the time axis for the 30-s data (in orange) was multiplicatively rescaled. Superposition between the two curves was indexed by eta-square (η2) indicated in the bottom right of each graph, the highest values being highlighted in red. B) Local field potential (LFP) signal power in the mPFC (medial prefrontal cortex), in the theta (5-15Hz) and gamma (40-80Hz) bands. Left side, raw data: theta and gamma power was expressed as percent change from the baseline (BL), and represented from odor onset (green vertical line) to shock arrival which occurs either at 20s (Group 20s, in blue) or at 30s (Group 30s, in orange). Right side, rescaled curves. The red asterisk in the upper left corner of a graph signals a significant time x interval duration interaction. Open circles on each curve signal the values that are significantly different from the baseline (p<0.05). Superposition between the two curves was indexed by eta-square (η2) indicated in the bottom right of each graph, the highest values being highlighted in red.

Figure 7B, upper left part, presents the temporal curves obtained for theta activity power in mPFC in the two groups. The ANOVA carried out in the four recording sites revealed no effect of Group, but a significant effect of Time in mPFC (F_17,238_=2.44, p=0.002), and a significant Time×Group interaction in mPFC (F_17,238_=2.32, p=0.003), BLA (F_17,153_=2.18, p=0.007) and DMS (F_17,238_=2.11, p=0.01) (Supplementary Fig. S4, left part). When the time axis for the 30-s data was multiplicatively rescaled to assess scalar timing, η^2^ was greater under the multiplicative transform than under no rescaling in mPFC, BLA and DMS (Figure 7B upper right part, and Supplementary Fig. S4 right part).

Figure 7B, lower left part, presents the temporal curves obtained for gamma activity power in mPFC in the two groups. The ANOVA carried out in the four recording sites (Supplementary Fig. S5, left part) revealed no effect of Group, but a significant effect of Time in mPFC (F_17,238_=4.21, p<0.0001), PIR (F_17,221_=2.12, p=0.007) and DMS (F_17,238_=3.45, p<0.0001), and a significant Time×Group interaction in mPFC (F_17,221_=3.30, p<0.0001) and DMS (F_17,238_=2.45, p=0.001). When the time axis for the 30s data was multiplicatively rescaled to assess scalar timing, η^2^ was greater under the multiplicative transform than under no rescaling in mPFC and DMS (Figure 7B, lower right part, and Supplementary Fig. S5 right part) In summary, for the respiration and theta temporal curves, parallel training with either 20s or 30s odor-shock interval duration in different animals brought globally the same kind of data as when animals were trained successively to the two intervals. Concerning gamma activity, parallel training revealed a modulation of activity power temporal curves by the interval’s duration in mPFC and DMS in a manner that respected scalar property.

## DISCUSSION

The present study is the first to correlate brain neural dynamics associated with behavioral expression of interval timing at early stages of Pavlovian aversive conditioning. This was made possible thanks to the use of a sensitive behavioral index, respiration, which allowed getting a readout of the animal’s interval timing ability without training the animal in a long and complex operant behavioral task. Here we show that in a successive training paradigm where the initially learned odor-shock interval duration is shifted to a new one, animals exhibiting good timing performances as assessed through respiration rate, also show temporal patterns of theta activity power modulated by the interval duration in the four recording sites, and those measured in mPFC and DMS followed the scalar property. In addition, the temporal patterns of theta activity in the mPFC faithfully follow the temporal patterns of respiration. In contrast, temporal patterns of gamma activity power modulated by the interval duration were observed only in mPFC, and they failed to respect scalar property (see a summary of the data on Table 1, upper part).

**Table 1:** Summary of the data obtained for theta and gamma activity power in Good Timers animals in the two experimental paradigms (Successive training and parallel training) and in the four recording sites, for the criteria relevant to interval timing, ie significant Interval x Time interaction and respect of scalar property. mPFC: medial prefrontal cortex; BLA: basolateral amygdala; PIR: olfactory piriform cortex; DMS: dorso-medial striatum.

| Successive training | Theta |  | Gamma |  |
| --- | --- | --- | --- | --- |
|  | Interval x Time | Scalar property | Interval x Time | Scalar property |
| mPFC | x | x | x |  |
| BLA | x |  |  |  |
| PIR | x |  |  |  |
| DMS | x | x |  |  |

**Table 1:** Summary of the data obtained for theta and gamma activity power in Good Timers animals in the two experimental paradigms (Successive training and parallel training) and in the four recording sites, for the criteria relevant to interval timing, ie significant Interval x Time interaction and respect of scalar property.
| Parallel training | Theta |  | Gamma |  |
| --- | --- | --- | --- | --- |
|  | Interval x Time | Scalar property | Interval x Time | Scalar property |
| mPFC | x | x | x | x |
| BLA | x | x | x | x |
| PIR |  |  |  |  |
| DMS | x | x |  |  |

### Interval timing and level of stress

The present work highlights that respiratory rhythm can be used as a reliable index of interval timing providing that the level of stress of the animal is not too high. Indeed, we could distinguish two subgroups of animals based on their individual temporal pattern of respiratory frequency during the odor-shock interval, and its modification following the shift to a new interval duration. Good Timers presented temporal patterns that were modulated by the duration of the interval in a way that respected scalar property. In contrast, Bad Timers showed changes in respiratory frequency during the interval, but these changes were not modulated by the duration of the interval.

At the behavioral level, Bad Timers showed more USV emissions during the two training sessions. USV emission is known to narrow and lower the respiratory frequency range (Frysztak and Neafsey, 1991; Hegoburu et al, 2011; Dupin et al, 2019), which might have impeded the development of clear-cut temporal patterns. In the present study, all the animals except two (out of 30) produced USVs albeit at different rates. This is in contrast to what is usually observed in non-implanted animals, where the ratio of vocalizing animals is generally around 50% for the same foot-shock parameters (Wöhr et al, 2005; Hegoburu et al, 2011; Shionoya et al, 2013). The reason for this high rate of USV vocalizing animals in the present study might be related to their housing condition, as animals implanted with electrodes were housed individually. It has been shown that single housing increases anxiety and biological markers of stress as compared to social housing (Arakawa, 2018; Manouze et al, 2019) and the amount of USV emission is enhanced when the level of anxiety is increased (Wöhr et al, 2005). Indeed, more anxious rats (as assessed through the elevated plus maze) were shown to exhibit more freezing and to be more likely to vocalize than less anxious animals (Borta et al, 2006).

A few studies have addressed the impact of emotion on temporal behavior in animals. Among them, studies using a peak interval procedure (Brown et al, 2007; Meck and Macdonald, 2007; Garces et al 2018) or a temporal bisection task (Faure et al, 2013; Kamada and Hata, 2019) showed that the presentation of foot-shocks or of a cue with a negative valence through its association with foot-shock, produces a drastic disruption of the temporal behavior. In the present study, higher levels of stress in the Bad Timers subgroup might have induced the same effect on their timing performances.

It might be argued that the absence of interval-related temporal patterns in respiratory frequency in Bad Timers rats does not mean that these animals were not capable to time intervals, but rather that this ability could not be assessed through respiratory rhythm. Using another index like, for example, heart rate, might have allowed detecting interval timing in these rats (Sananes and Campbell, 1989). However, as neural activity power in Bad Timers, akin to their respiration rate, was not modulated by interval duration, we feel confident that our classification based on respiratory patterns does indeed reflect the animal’s timing ability. Nevertheless, it remains possible that other neural correlates of interval timing might have existed even in animals expressing no temporal behavioral pattern, as described by Tallot et al (2020).

Some authors have proposed that temporal relations between stimuli are automatically encoded, and are the foundation of associative learning (Balsam and Gallistel, 2009). Based on this assumption, it might be expected that animals with poor timing abilities should present lower learning performances than animals with good timing abilities. In the present study Bad Timers and Good Timers showed similar levels of learned odor fear during the 48h retention test. However, analyzing more in depth the fear memory strength, by investigating for example if the memory is more resistant to extinction or persists longer in Good Timers than in Bad Timers, might reveal learning deficits in Bad Timers that were not detected here.

### Changes in theta and gamma activity power

Importantly, the temporal patterns of LFP signal power in the theta band were clearly different between Good Timers and Bad Timers subgroups suggesting that this group subdivision initially based on respiration temporal patterns is also associated with differences in neural processes. In the Good Timers subgroup, the temporal patterns of theta activity power were modulated by the duration of the odor-shock interval in the four recording sites, and respected scalar property in mPFC and DMS. In contrast, in the Bad Timers subgroup, theta activity power did not change compared to baseline during the odor-shock interval. Concerning activity in the gamma band, although signal power increased in Good Timers and Bad Timers during the interval, only temporal patterns in the mPFC in the Good Timers subgroup were modulated by the duration of the odor-shock interval, but without following scalar property. These data suggest that the learning of interval duration in our paradigm might be supported preferentially by theta oscillatory activity in a network including mainly the mPFC and DMS. These observations are partly in line with the data reported by Dallérac et al (2017). In that study, interval timing ability was assessed using a conditioned suppression paradigm, in which a learned aversive cue suppresses ongoing operant behavior consisting in lever pressing for food. Aversive conditioning consisted of a tone paired with a mild electric footshock delivered 30s after tone onset. After several weeks of training, rats showed a bell-shaped curve of lever-pressing suppression, the maximum conditioned suppression being at a time close to the shock arrival. Furthermore, shifting the tone-shock interval from 30s to 10s yielded a rapid shift in the peak of suppression. The authors examined oscillatory activity power in DMS and BLA and reported that the power of both theta and gamma oscillations increased during the tone with a maximum that shifted as a result of a change in tone-shock interval duration. Moreover, a significant increase in coherence between these structures was found in the low theta band. This led the authors to propose that learning of the association *per se* is modulated by the amygdalo-striatal projection through gamma oscillations while the timing component of the association would be driven by theta oscillations.

### Successive versus parallel training with different interval durations

In the present study, most of the animals were trained successively to the two interval durations. Such a paradigm allows comparing changes within animals, thus discounting between animals’ differences, particularly concerning LFP signal power that can be very different between animals due to differences in recording electrode tip position. However, the data obtained during the Shift session might have been partly confounded by the Conditioning session. Indeed, during the Shift session, the animals not only have to learn the new duration but also to forget the previous one. We therefore compared animals in parallel groups, using a single training session with either 20s or 30s odor-shock interval duration. The data were similar to those obtained with successive training for respiration temporal patterns, allowing us to assume that these patterns are mostly linked to interval timing per se and not to training levels. Concerning oscillatory activity data, although both parallel and successive training led to temporal patterns modulated by interval duration, these patterns were slightly different between the two conditions particularly concerning gamma activity (see a summary of the data on Table 1, lower part). More specifically, in animals trained in parallel sessions, temporal patterns of gamma activity power follow scalar property in mPFC and BLA while they do not in animals trained in successive sessions. This suggests that gamma activity in these structures keeps a trace of the previously learned interval duration, which could have blurred the 30s temporal pattern in our successive paradigm, thus preventing the scalar property to be observed.

### Does respiratory rhythm modulate the temporal patterns observed in theta band?

In a recent study, Tallot et al (2020) using a discriminative fear conditioning paradigm where animals had to discriminate between a CS+ (associated with a footshock) and a CS− (never associated with a footshock), showed that, early in training, dynamics of neuronal oscillations in an amygdalo– prefronto–striatal network are modified during the CS+ in a manner related to the CS–US time interval. Interestingly these changes occurred while the behavioral response assessed via freezing was similar for CS+ and CS-, indicating high level of fear generalization, thus precluding any causal relationship between time-related modulations in LFPs, and freezing behavior. In the present study, the monitoring of respiration in addition to freezing allowed us to correlate changes in respiratory rate with changes in oscillatory activity power. We found that the temporal patterns of respiratory frequency and theta activity power present striking similarities. Indeed the time of the respiratory frequency peak (which occurs around 4Hz) was positively correlated with that of the theta activity power peak. In addition, when the duration of the interval was changed, there was a positive correlation between the shift in the respiratory frequency peak time and that of the theta activity power in mPFC. This suggests a potential role for respiratory rhythm in the emergence of theta temporal patterns in the mPFC. Indeed, several recent studies in the literature have shown that the respiratory rhythm in addition to its impact on olfactory regions (for a review, see Buonviso et al., 2006), also entrains oscillations in widespread brain regions including those involved in the fear network like the mPFC and amygdala (for a review, see Tort et al., 2018; Heck et al, 2017). This suggests that the breathing rhythm, akin to slow oscillatory rhythms, could help coordinate neural activity across distant brain regions (Jensen and Colgin, 2007; Heck et al., 2017), and potentially modulate cognitive processes. In the present study, it was not possible to analyze the temporal pattern of activity in the 3-5 Hz delta band due to edge effects due to shock artefact. However, others have found that a 4Hz oscillation develops in the mPFC during freezing behavior and that this oscillation was driven by respiration (Moberly et al, 2016; Karalis et al, 2016; Bagur and Benchenane, 2018). In line with these studies, we recently showed that respiratory frequency and delta activity frequency were highly correlated during freezing (Dupin et al, 2019). It could be hypothesized that the respiration-triggered 4Hz oscillation, when it occurs, is particularly well suited to modulate the amplitude of theta and higher frequency oscillations. Importantly we previously reported that the emission of USV, that drastically lowers the respiratory rate down to 0.5-1.5Hz, disrupted the correlation between respiration and delta rhythm and was associated with a decrease in theta activity power (Dupin et al, 2019). This could explain why in Bad Timers animals, which present higher USV rates, theta activity power did not increase during the odor-shock interval.

In conclusion, the present work describes for the first time to our knowledge, some of the brain neural dynamics associated with behavioral expression of interval timing at early stages of Pavlovian aversive conditioning. Our study highlights the potential preferential role of theta rhythm in the learning of interval durations, particularly in mPFC and DMS. In addition, our data suggest the novel idea that the respiratory rhythm might take an active part in the setting of theta activity dynamics, at least in the mPFC. Whether this role is restricted to the use of an olfactory conditioned stimulus that is sampled through active sniffing or can be observed for another sensory modality, will deserve further investigation. Finally, we were able to show that temporal patterns of theta activity develop concomitantly to temporal patterns of respiration, in a single training session, thus bringing support to the proposal that learning the association between two stimuli co-occurs with the learning of their temporal relationships (Balsam and Gallistel, 2009).

## METHODS

### Animals

Data were obtained from thirty male Long Evans rats (250-270g at their arrival, Janvier Labs, France). They were housed individually at 23°C and maintained under a 12 h light–dark cycle (lights on from 8:00 am to 8:00 pm). Food and water were available ad libitum. All experiments and surgical procedures were conducted in strict accordance with the European Community Council Directive of 22nd September 2010 (2010/63/UE) and the national ethics committee (APAFIS#10606). Care was taken at all stages to minimize stress and discomfort to the animals. Some of the behavioral and electrophysiological data collected on these animals have been used for another paper (Dupin et al, 2019).

### Surgery

Animals were anesthetized with Equithesin, a mixture of chloral hydrate (127 mg/kg, i.p.) and sodium pentobarbital (30 mg/kg, i.p.), and placed in a stereotaxic frame (Narishige, Japan) in a flat skull position. The level of anesthesia was held constant with regular injections of Equithesin throughout the surgery. Monopolar stainless steel recording electrodes (100 µm in diameter) were then stereotaxically implanted in the left hemisphere in the four brain areas: mPFC (Prelimbic part, AP: +3.0mm; L: +0.8mm; DV: −3.5mm), DMS (AP: +1.2mm, L: +2.2mm, DV: −3.5mm), PIR (AP: −1.8, L: +5.5mm, DV: −8 mm) and BLA (AP : −2.8mm, L : +4.9mm, DV : −7.5mm). Accurate positioning in the PIR was achieved using the characteristic profiles of evoked field potential induced in the PIR in response to electrical stimulation of the olfactory bulb (Haberly, 1973). For this, a bipolar stimulation electrode (made of two 100-μm stainless-steel wires with a tip separation of 500 μm) was lowered transiently in the olfactory bulb to facilitate positioning in the PIR and withdrawn thereafter. A screw inserted in the skull bone above the right parietal lobe served as a reference and ground electrode. The four recording electrodes and the ground electrode were connected to a telemetry transmitter (rodentPACK system, EMKA Technologies, Paris, France) fixed to the rat’s skull surface by dental acrylic cement and anchored with a surgical screw placed in the frontal bone. The animals were allowed to recover for two weeks following surgery.

### Experimental apparatus

The apparatus has been described in detail in a previous study (Hegoburu et al., 2011). It consisted of a whole body customized plethysmograph (diameter 20 cm, height 30 cm, Emka Technologies, France) placed in a sound-attenuating cage (L 60 cm, W 60 cm, H 70 cm, 56dB background noise). The plethysmograph was used to measure respiratory parameters in behaving animals. The bottom of the animal chamber was equipped with a shock floor connected to a programmable Coulbourn shocker (Bilaney Consultants GmbH, Düsseldorf, Germany). Three Tygon tubing connected to a programmable custom olfactometer were inserted in the tower on the top of the plethysmograph to deliver air and odorants. Deodorized air flowed constantly through the cage (2 L/min). When programmed, an odor (McCormick Pure Peppermint; 2 L/min; 1:10 peppermint vapor to air) was introduced smoothly in the air stream through the switching of a solenoid valve (Fluid automation systems, CH-1290 Versoix, Switzerland), thus minimizing its effect on change in pressure. A ventilation pump connected to a port at the bottom of the animal chamber allowed drawing air out of the plethysmograph (at a rate of up to 2 L/min), thus maintaining a constant airflow that did not interact with the animal’s breathing pattern. The experimental setup also allowed recording two additional behavioral responses from the animals: freezing behavior was monitored via two video cameras fixed on the walls of the sound-attenuating cage and ultrasonic vocalizations (USV) were recorded via a condenser ultrasound microphone (Avisoft-Bioacoustics CM16/CMPA, Berlin, Germany) inserted in a tower adapted to the ceiling of the plethysmograph.

### Fear conditioning paradigm (Figure 1A)

After the recovery period, the animals were handled individually and placed in the experimental apparatus for 30 min per day during 3 to 4 days before the beginning of the experiments in order to familiarize them with being manipulated and connected to the telemetry transmitter.

For the Conditioning session, the telemetry transmitter was plugged onto the animal’s head and the rat was allowed free exploration during the first 4 min, then an odor was introduced into the plethysmograph cage for 20s (Group 20s, n=21), the last second of which overlapped with the delivery of a 0.4mA foot-shock (1s). The animal received ten odor-shock trials, with an intertrial interval of 4 min. After the last pairing, the transmitter was unplugged, and the animal returned to its home cage. In a small group of animals, the odor-shock interval was set at 30s (Group 30s, n=9), while everything else was kept the same as for Group 20s.

The conditioned fear memory was assessed in group 20s animals during a Retention test carried out 48h after conditioning. For the Retention test, the rat was placed in the experimental cage (equipped with new visual cues and with a plastic floor to avoid contextual fear expression) and allowed a 4-min odor-free period. The CS odor was then presented five times for 20s with a 4-min intertrial interval. Two weeks later, these animals were submitted to a Shift session during which the initial 20s-odor-shock interval was shifted to 30s. The animal was connected to the telemetry recording system and received ten 30s-odor-shock trials. After the last pairing the transmitter was unplugged, and the animal returned to its home cage.

### Data acquisition and preprocessing

The respiratory signal collected from the plethysmograph was amplified and sent to an acquisition card (MC-1608FS, Measurement Computing, USA; Sampling rate = 1000 Hz) for storage and offline analysis. The detection of the respiratory cycles was achieved using an algorithm described in a previous study (Roux et al., 2006). This algorithm performs two main operations: signal smoothing for noise reduction, and detection of zero-crossing points to define accurately the inspiration and expiration phase starting points. Momentary respiratory frequency was determined as the inverse of the respiratory cycle (inspiration plus expiration) duration.

The video signals collected through the two cameras were transmitted to a video acquisition card and a homemade acquisition software. Offline, freezing behavior defined as the absence of any visible movement except that due to breathing (Blanchard and Blanchard, 1969), was automatically detected using a Labview homemade software and further verified by an experimenter (for a detailed description, see Dupin et al, 2019).

For USV recording, the ultrasound microphone was connected to a recording interface (UltraSoundGate 116 Hb, Avisoft-Bioacoustics, Berlin, Germany) with the following settings: sampling rate = 214285 Hz; format = 16 bit (Wohr et al., 2005). Recordings were transferred to Avisoft SASLab Pro (version 4.2, Avisoft Bioacoustics, Berlin, Germany) and a Fast Fourier Transform (FFT) was conducted. Spectrograms were generated with an FFT-length of 512 points and a time window overlap of 87.5% (100% Frame, FlatTop window). These parameters produced a spectrogram at a frequency resolution of 419 Hz and a time resolution of 0.29 ms. The acoustic signal detection was provided by an automatic whistle tracking algorithm with a threshold of −20 dB, a minimum duration of 0.01 s and a hold time of 0.02 s. However, the accuracy of detection was verified trial by trial by an experienced user. The main parameters used in the present study were extracted using Avisoft SASLab Pro and concerned the duration as well as the start and end time of USV calls. No band pass filter has been applied during USV recording.

Local field potentials (LFP) were collected by telemetry via a four-channel wireless miniature transmitter (<5.2g, RodentPack EMKA Technology). LFP signals were amplified (x1000), filtered (between 0.1 and 100 Hz), digitized (sampling frequency: 1000 Hz) and stored on a computer for offline analysis. The LFP signals were first individually inspected in order to eliminate artifacts due to signal saturation or transient signal loss. The selection was made for each recording site separately and proceeded as follows: when the duration of an artifact exceeded 5s over the recording period analyzed, the trial was excluded. When the number of excluded trials exceeded 5 (out of the 10 trials) then the recording site was excluded for this animal. After the elimination of artifacted signals, the number of animals per recording site were the following: BLA, n=17; CPF, n=20; PIR, n=20; DMS, n=19 for Group 20s; BLA, n=6; CPF, n=8; PIR, n=8; DMS, n=9 for Group 30s.

### Data analysis (Figure 1B)

The different data (respiration, USV, behavior, LFP signals) were synchronized offline via a TTL synchronization signal generated at the beginning of each training session. Once synchronized, the data were analyzed using custom-written scripts in Python.

The temporal course of respiratory frequency and LFP signal power during the 20s (or 30s) odor-shock interval was assessed during the Conditioning and Shift sessions as follows. For each animal, the instant respiratory frequency was averaged second by second for each trial of the training session. The power spectral density (PSD) of the LFP signals was calculated using the continuous Morlet wavelet transform (Kronland-Martinet et al., 1987). The Morlet wavelet estimated the amplitude of the signal at each time and frequency bin. Two frequency bands were chosen for the subsequent analyses: theta (5-15Hz) and gamma (40-80Hz), and the mean power in each band was calculated second by second. The resulting values were normalized to baseline defined as the average power over the 20s preceding odor onset. The resulting curves of each trial (for respiratory frequency or LFP PSD) were averaged across the trials 2 to 9 of the session (Trial 1 was not included because the animal had not yet been exposed to the Odor-shock association) for a given animal and then averaged among animals of the same experimental group.

During the Conditioning and Shift sessions, the amount of Freezing and USV emission during the Odor-Shock intervals was quantified. Freezing rate was expressed as a percentage of the sampled period total duration, averaged across the 9 trials for a given animal and then averaged among animals of the same experimental group. In addition, for each animal, the number of USV emitted during the Odor-Shock intervals throughout the whole session was measured and averaged among animals of the same experimental group.

During the Retention test, the fear memory strength was assessed by measuring the duration of the animal’s freezing response before introduction of the first odor, and during each odor presentation. The obtained values were then expressed as a percentage of the sampled period total duration, averaged across the trials for a given animal, and averaged among animals of the same experimental group.

### Statistical analysis

All analyses were performed with Systat 13.0® software. For each test, the significance level was set at p ≤ 0.05.

In Group 20s during training, the per-second temporal curves of respiratory frequency and LFP signal power observed during the Odor-shock interval for the two interval durations were compared using a two-way ANOVA with Interval duration (20s or 30s) and Time (1s to 18s) as repeated measures factors. Post-hoc pairwise comparisons were then carried out when allowed by the ANOVA results.

In Group 20s during Conditioning and Shift sessions, the mean global freezing rate and USV number measured during the Odor-shock interval for the two interval durations were compared using a two-way ANOVA with Subgroup (Good Timers or Bad Timers, see Results, first paragraph, for Subgroup definition) as an independent factor and Interval duration (20s or 30s) as a repeated measures factor. Posthoc pairwise comparisons were then carried out when allowed by the ANOVA results.

In Group 20s during the Retention test, the learned fear response to the odor was analyzed using a two-way ANOVA with Group (Good Timers or Bad Timers) as an independent factor and Period (Pre-odor vs Odor) as a repeated measures factor. Posthoc pairwise comparisons were then carried out when allowed by the ANOVA results.

The per-second temporal curves of respiratory frequency and LFP signal power observed during the first training session in Group 20s and Group 30s were compared using a two-way ANOVA with Interval duration (20s or 30s) as an independent factor and Time (1s to 18s) as a repeated measures factor. Posthoc pairwise comparisons were then carried out when allowed by the ANOVA results.

### Histology

At the end of the experiment, the animals were injected intraperitoneally with a lethal dose of pentobarbital, their brains were removed, postfixed, cryoprotected in sucrose (20%). The brains were then sectioned (40µm coronal slices) for verification of electrodes tips by light microscopy. Areas targeted by the electrodes in the four implanted brain regions have been reported on brain atlas coronal sections (Supplementary Fig. S6).

## ACKNOWLEDGEMENTS

This work was supported by the Centre National de la Recherche Scientifique, the LABEX CORTEX (ANR-11-LABX-0042) of Université de Lyon, within the program “Investissements d’Avenir” (ANR-11-IDEX-0007) operated by the French National Research Agency, and Partner University Funds “Emotion & Time”.

We thank Julie Boulanger-Bertolus for valuable discussions of the data and careful reading of the manuscript, Marc Thevenet for technical assistance, and Ounsa Ben-Hellal for taking care of the animals.

Correspondence should be addressed to Maryne Dupin or Anne-Marie Mouly

## AUTHOR CONTRIBUTIONS

M.D. and A.-M.M. designed the study; M.D. did the experiments; B.M. helped with the design of the experimental setup; M.D., S.G. and A.-M.M. carried out the analyses; M.D. and A.-M.M. wrote the manuscript; V.D. discussed the data, commented and edited the manuscript.

## REFERENCES

Arakawa H (2018) Ethological approach to social isolation effects in behavioral studies of laboratory rodents. Behavioural brain research 341:98–108.

Arcediano F, Escobar M, Miller RR (2003) Temporal integration and temporal backward associations in human and nonhuman subjects. Learning & behavior 31:242–256.

Bagur S, Lefort JM, Lacroix MM, de Lavilléon G, Herry C, Billand C, Geoffroy H, Benchenane K (2018) Dissociation of fear initiation and maintenance by breathing-driven prefrontal oscillations. bioRxiv 468264; doi: https://doi.org/10.1101/468264

Balsam PD, Gallistel CR (2009) Temporal maps and informativeness in associative learning. Trends in neurosciences 32:73–78.

Blanchard RJ, Blanchard DC (1969) Crouching as an index of fear. Journal of comparative and physiological psychology 67:370–375.

Borta A, Wohr M, Schwarting RK (2006) Rat ultrasonic vocalization in aversively motivated situations and the role of individual differences in anxiety-related behavior. Behavioural brain research 166:271–280.

Boulanger-Bertolus J, Hegoburu C, Ahers JL, Londen E, Rousselot J, Szyba K, Thevenet M, Sullivan-Wilson TA, Doyere V, Sullivan RM, Mouly AM (2014) Infant rats can learn time intervals before the maturation of the striatum: evidence from odor fear conditioning. Frontiers in behavioral neuroscience 8:176.

Brown BL, Hemmes NS, Cabeza de Vaca S (1992) Effects of intratrial stimulus change on fixed interval performance: the roles of clock and memory processes. Animal Learning and Behaviour 20:83–93.

Brown BL, Richer P, Doyere V (2007) The effect of an intruded event on peak-interval timing in rats: isolation of a postcue effect. Behavioural processes 74:300–310.

Bueti D, Macaluso E (2010) Auditory temporal expectations modulate activity in visual cortex. NeuroImage 51:1168–1183.

Buhusi CV, Meck WH (2005) What makes us tick? Functional and neural mechanisms of interval timing. Nature reviews Neuroscience 6:755–765.

Buonviso N, Amat C, Litaudon P (2006) Respiratory modulation of olfactory neurons in the rodent brain. Chemical senses 31:145–154.

Clugnet MC, Price JL (1987) Olfactory input to the prefrontal cortex in the rat. Ann NY Acad Sci 510:231–235.

Corcoran KA, Quirk GJ (2007) Activity in prelimbic cortex is necessary for the expression of learned, but not innate, fears. The Journal of neuroscience: the official journal of the Society for Neuroscience 27:840–844.

Coull JT, Cheng RK, Meck WH (2011) Neuroanatomical and neurochemical substrates of timing. Neuropsychopharmacology: official publication of the American College of Neuropsychopharmacology 36:3–25.

Dallerac G, Graupner M, Knippenberg J, Martinez RC, Tavares TF, Tallot L, El Massioui N, Verschueren A, Hohn S, Bertolus JB, Reyes A, LeDoux JE, Schafe GE, Diaz-Mataix L, Doyere V (2017) Updating temporal expectancy of an aversive event engages striatal plasticity under amygdala control. Nature communications 8:13920.

Datiche F, Cattarelli M (1996) Reciprocal and topographic connections between the piriform and prefrontal cortices in the rat: a tracing study using the B subunit of the cholera toxin. Brain research bulletin 41:391–398.

Davis M, Schlesinger LS, Sorenson CA (1989) Temporal specificity of fear conditioning: effects of different conditioned stimulus-unconditioned stimulus intervals on the fear-potentiated startle effect. Journal of experimental psychology Animal behavior processes 15:295–310.

Diaz-Mataix L, Tallot L, Doyere V (2014) The amygdala: a potential player in timing CS-US intervals. Behavioural processes 101:112–122.

Diaz-Mataix L, Ruiz Martinez RC, Schafe GE, LeDoux JE, Doyere V (2013) Detection of a temporal error triggers reconsolidation of amygdala-dependent memories. Current biology : CB 23:467–472.

Dupin M, Garcia S, Boulanger-Bertolus J, Buonviso N, Mouly AM (2019) New Insights from 22-kHz Ultrasonic Vocalizations to Characterize Fear Responses: Relationship with Respiration and Brain Oscillatory Dynamics. eNeuro 6.

Faure A, Es-Seddiqi M, Brown BL, Nguyen HP, Riess O, von Horsten S, Le Blanc P, Desvignes N, Bozon B, El Massioui N, Doyere V (2013) Modified impact of emotion on temporal discrimination in a transgenic rat model of Huntington disease. Frontiers in behavioral neuroscience 7:130.

Frysztak RJ, Neafsey EJ (1991) The effect of medial frontal cortex lesions on respiration, “freezing,” and ultrasonic vocalizations during conditioned emotional responses in rats. Cerebral cortex 1:418–425.

Garces D, El Massioui N, Lamirault C, Riess O, Nguyen HP, Brown BL, Doyere V (2018) The Alteration of Emotion Regulation Precedes the Deficits in Interval Timing in the BACHD Rat Model for Huntington Disease. Frontiers in integrative neuroscience 12:14.

Headley DB, Weinberger NM (2011) Gamma-band activation predicts both associative memory and cortical plasticity. The Journal of Neuroscience: the official journal of the Society for Neuroscience 31:12748–12758.

Heck DH, McAfee SS, Liu Y, Babajani-Feremi A, Rezaie R, Freeman WJ, Wheless JW, Papanicolaou AC, Ruszinko M, Sokolov Y, Kozma R (2016) Breathing as a Fundamental Rhythm of Brain Function. Frontiers in neural circuits 10:115.

Hegoburu C, Sevelinges Y, Thevenet M, Gervais R, Parrot S, Mouly AM (2009) Differential dynamics of amino acid release in the amygdala and olfactory cortex during odor fear acquisition as revealed with simultaneous high temporal resolution microdialysis. Learning & memory 16:687–697.

Hegoburu C, Shionoya K, Garcia S, Messaoudi B, Thevenet M, Mouly AM (2011) The RUB Cage: Respiration-Ultrasonic Vocalizations-Behavior Acquisition Setup for Assessing Emotional Memory in Rats. Frontiers in behavioral neuroscience 5:25.

Hegoburu C, Parrot S, Ferreira G, Mouly AM (2014) Differential involvement of amygdala and cortical NMDA receptors activation upon encoding in odor fear memory. Learn Mem 21:651–655.

Jensen O, Colgin LL (2007) Cross-frequency coupling between neuronal oscillations. Trends in cognitive sciences 11:267–269.

Kamada T, Hata T (2019) Basolateral amygdala inactivation eliminates fear-induced underestimation of time in a temporal bisection task. Behavioural brain research 356:227–235.

Karalis N, Dejean C, Chaudun F, Khoder S, Rozeske RR, Wurtz H, Bagur S, Benchenane K, Sirota A, Courtin J, Herry C (2016) 4-Hz oscillations synchronize prefrontal-amygdala circuits during fear behavior. Nat Neurosci 19:605–612.

Kronland-Martinet R, Morlet J, Grossmann A (1987) Analysis of sound patterns through wavelet transforms. Int J Pattern Recog Art Intel 1:273–302.

LeDoux JE (2000) Emotion circuits in the brain. Annual review of neuroscience 23:155–184.

Manouze H, Ghestem A, Poillerat V, Bennis M, Ba-M’hamed S, Benoliel JJ, Becker C, Bernard C (2019) Effects of Single Cage Housing on Stress, Cognitive, and Seizure Parameters in the Rat and Mouse Pilocarpine Models of Epilepsy. eNeuro 6.

Matell MS, Meck WH (2004) Cortico-striatal circuits and interval timing: coincidence detection of oscillatory processes. Brain research Cognitive brain research 21:139–170.

McDonald AJ (1998) Cortical pathways to the mammalian amygdala. Progress in neurobiology 55:257–332.

McGeorge AJ, Faull RL (1989) The organization of the projection from the cerebral cortex to the striatum in the rat. Neuroscience 29:503–537.

Meck WH, Macdonald CJ (2007) Amygdala inactivation reverses fear’s ability to impair divided attention and make time stand still. Behavioral neuroscience 121:707–720.

Moberly AH, Schreck M, Bhattarai JP, Zweifel LS, Luo W, Ma M (2018) Olfactory inputs modulate respiration-related rhythmic activity in the prefrontal cortex and freezing behavior. Nat Commun 9:1528.

Parker KL, Chen KH, Kingyon JR, Cavanagh JF, Narayanan NS (2014) D1-dependent 4 Hz oscillations and ramping activity in rodent medial frontal cortex during interval timing. The Journal of neuroscience : the official journal of the Society for Neuroscience 34:16774–16783.

Paxinos G, Watson C (2007). The rat brain in stereotaxic coordinates. San Diego, CA: AcademicPress.

Popescu AT, Popa D, Pare D (2009) Coherent gamma oscillations couple the amygdala and striatum during learning. Nature neuroscience 12:801–807.

Roberts S (1981) Isolation of an internal clock. Journal of experimental psychology Animal behavior processes 7:242–268.

Roux SG, Garcia S, Bertrand B, Cenier T, Vigouroux M, Buonviso N, Litaudon P (2006) Respiratory cycle as time basis: an improved method for averaging olfactory neural events. Journal of neuroscience methods 152:173–178.

Sacco T, Sacchetti B (2010) Role of secondary sensory cortices in emotional memory storage and retrieval in rats. Science 329:649–656.

Sananes CB, Campbell BA (1989) Role of the central nucleus of the amygdala in olfactory heart rate conditioning. Behavioral neuroscience 103:519–525.

Sevelinges Y, Gervais R, Messaoudi B, Granjon L, Mouly AM (2004) Olfactory fear conditioning induces field potential potentiation in rat olfactory cortex and amygdala. Learn Mem 11:761–769.

Shionoya K, Hegoburu C, Brown BL, Sullivan RM, Doyere V, Mouly AM (2013) It’s time to fear! Interval timing in odor fear conditioning in rats. Frontiers in behavioral neuroscience 7:128.

Shuler MG, Bear MF (2006) Reward timing in the primary visual cortex. Science 311:1606–1609.

Tallot L, Doyère V (2020) Neural encoding of time in the animal brain. Neuroscience and Biobehavioral Reviews 115:146–163.

Tallot L, Graupner M, Diaz-Mataix L, Doyère V (2020) Beyond freezing: Temporal expectancy of an aversive event engages the amydgalo-prefronto-dorsostriatal network. Cerebral Cortex.

Tort ABL, Brankack J, Draguhn A (2018) Respiration-Entrained Brain Rhythms Are Global but Often Overlooked. Trends in neurosciences 41:186–197.

Wohr M, Borta A, Schwarting RK (2005) Overt behavior and ultrasonic vocalization in a fear conditioning paradigm: a dose-response study in the rat. Neurobiology of learning and memory 84:228–240.

Zelano C, Jiang H, Zhou G, Arora N, Schuele S, Rosenow J, Gottfried JA (2016) Nasal Respiration Entrains Human Limbic Oscillations and Modulates Cognitive Function. The Journal of Neuroscience: the official journal of the Society for Neuroscience 36:12448–12467.

