## Supplemental data for "Respiration and brain neural dynamics associated with interval timing during odor fear learning in rats"

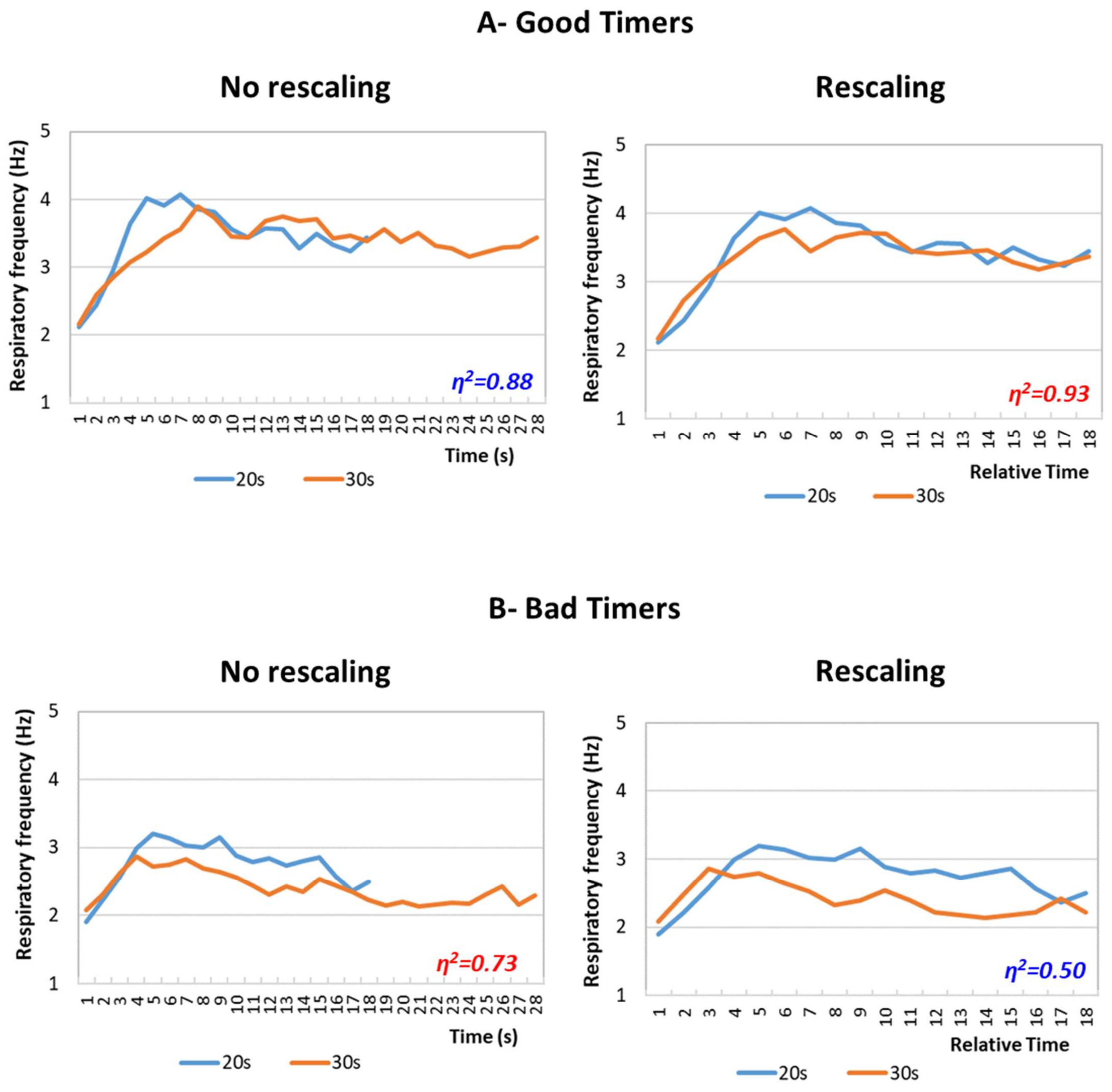


**Supplementary Figure S1:** *Rescaling of respiratory frequency curves during the odor-shock interval. A- Good Timers subgroup. Left side, raw data: respiratory frequency is represented from odor onset during the 20s (in blue) or the 30s (in orange) interval. Right side, rescaled curves: the time axis for the 30-s data (in orange) was multiplicatively rescaled. Superposition between the two curves was indexed by eta-square (η2) indicated in the bottom right of each graph, the highest values being highlighted in red. B- Bad Timers subgroup. Same explanations as in A.*

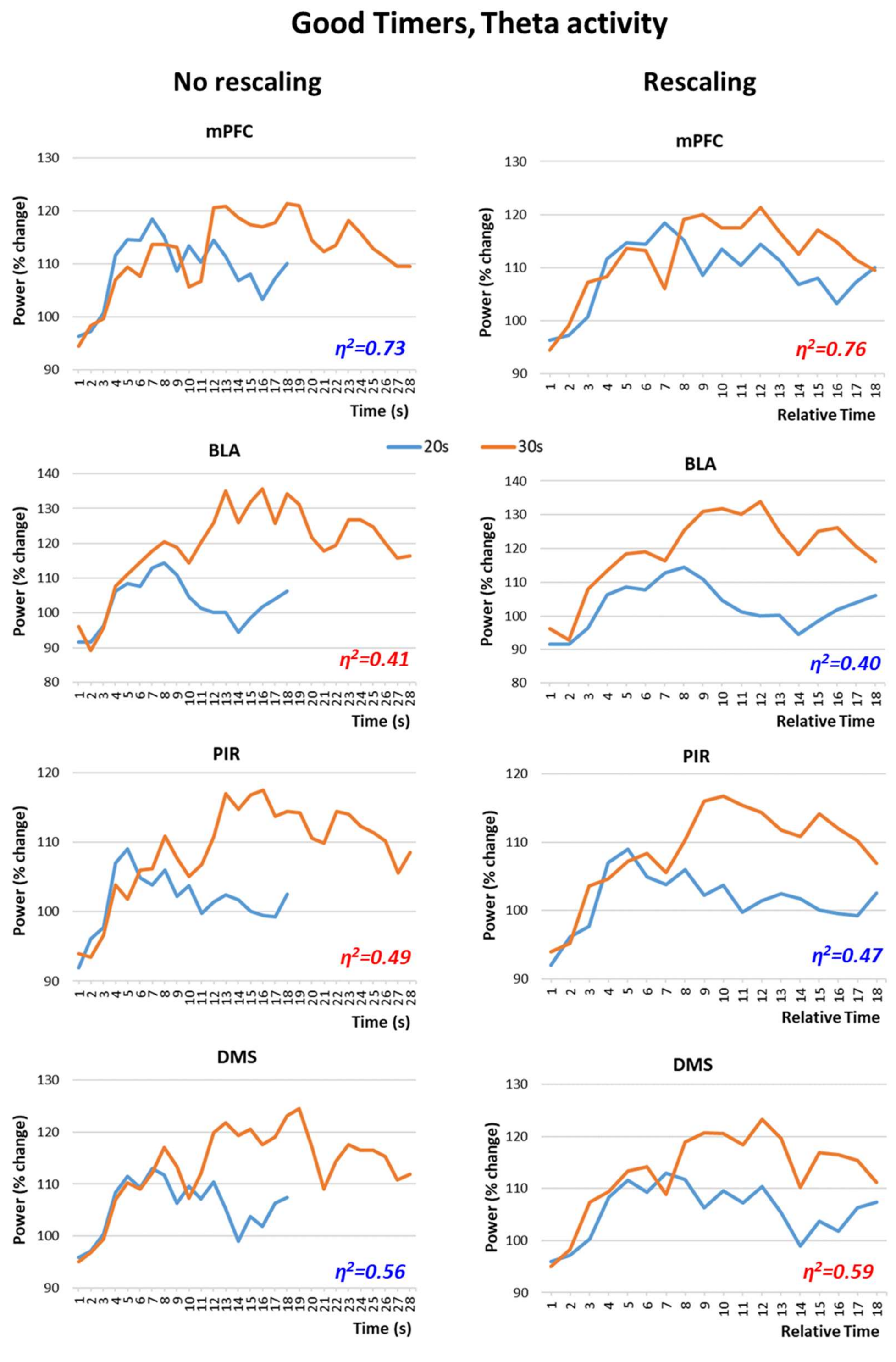


**Supplementary Figure S2:** *Rescaling of theta (5-15Hz) band mean power variations during the odor-shock interval in the Good Timers subgroup in the four recording sites. mPFC: medial prefrontal cortex; BLA: basolateral amygdala; PIR: olfactory piriform cortex; DMS: dorso-medial striatum. Left side, raw data: theta power changes are represented from odor onset during the 20s (in blue) or the 30s (in orange) interval. Right side, rescaled curves: the time axis for the 30-s data (in orange) was multiplicatively rescaled. Superposition between the two curves was indexed by eta-square ($\eta^2$) indicated in the bottom right of each graph, the highest values being highlighted in red.*

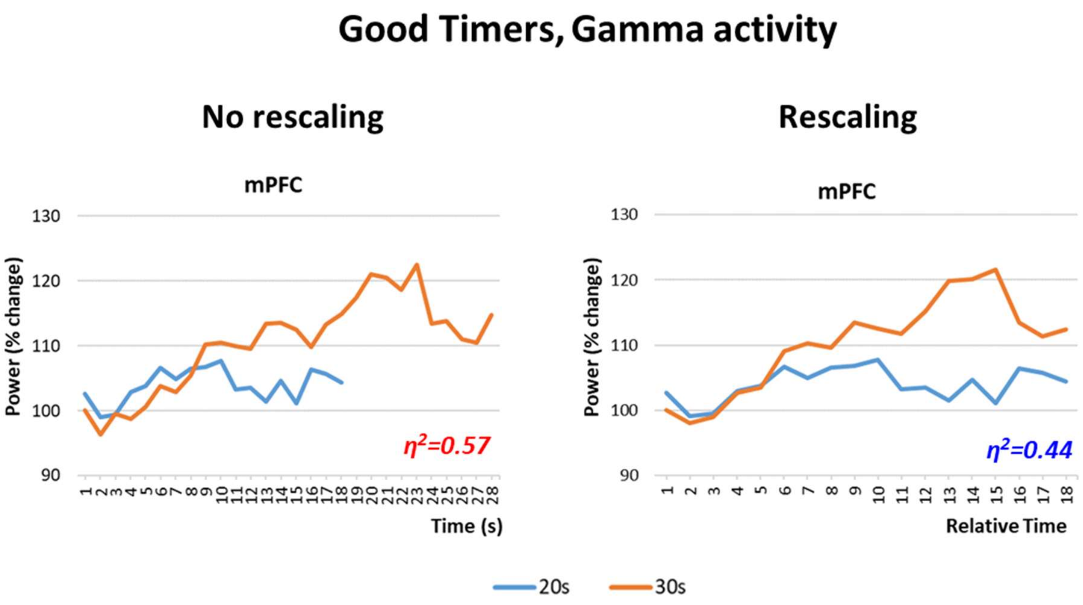


**Supplementary Figure S3:** *Rescaling of gamma (40-80Hz) band mean power variations during the odor-shock interval in the Good Timers subgroup in the mPFC: medial prefrontal cortex. Left side, raw data: gamma power changes are represented from odor onset during the 20s (in blue) or the 30s (in orange) interval. Right side, rescaled curves: the time axis for the 30-s data (in orange) was multiplicatively rescaled. Superposition between the two curves was indexed by eta-square ($\eta^2$) indicated in the bottom right of each graph, the highest values being highlighted in red.*

### Parallel training, Theta activity

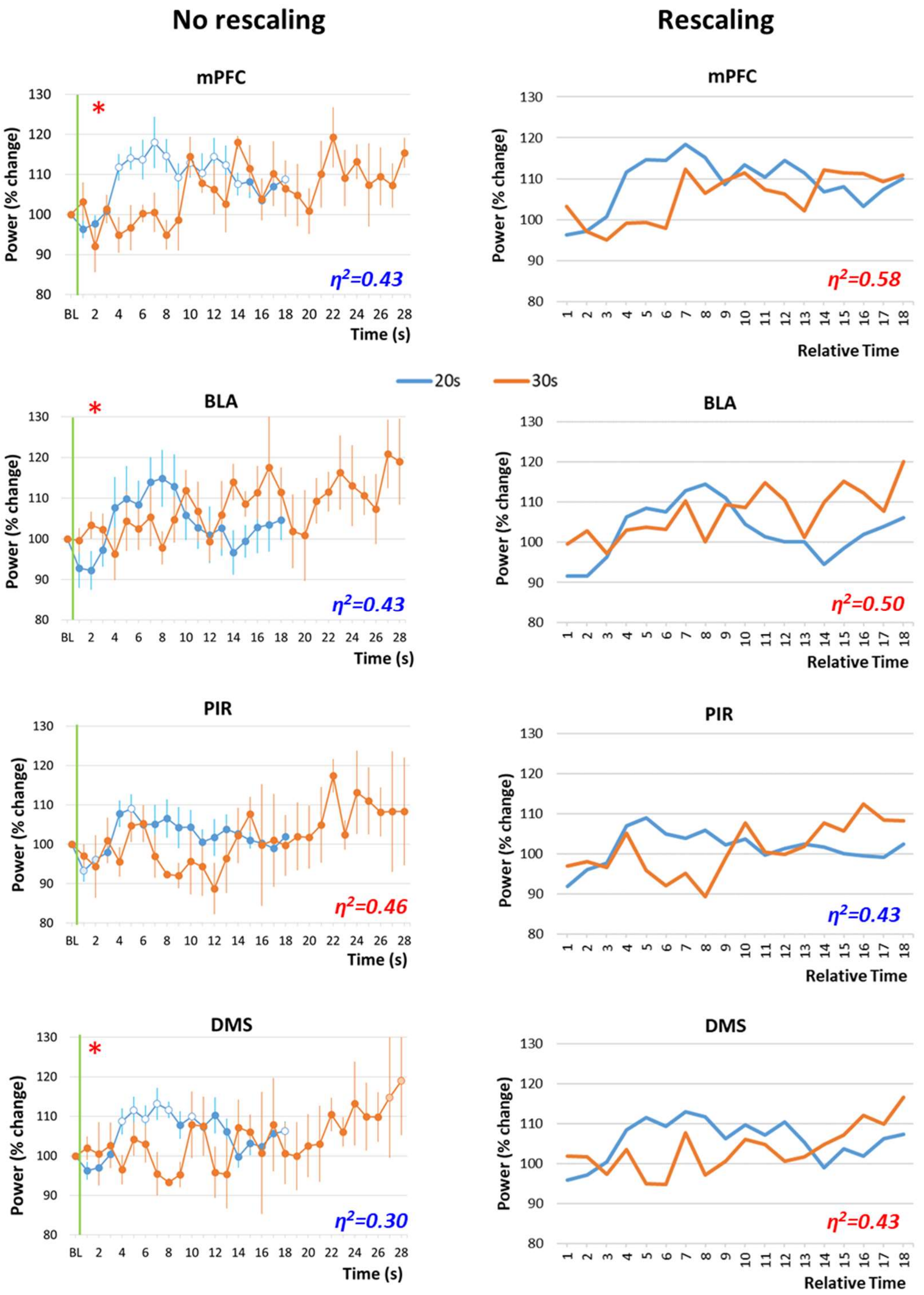


**Supplementary Figure S4:** *Theta (5-15Hz) band mean power variations in Group 20s and Group 30s, during the odor-shock interval in the four recording sites. mPFC: medial prefrontal cortex; BLA: basolateral amygdala; PIR: olfactory piriform cortex; DMS: dorso-medial striatum. Left side, raw data: theta power changes are represented from odor onset (green vertical line) during the 20s (in blue) or the 30s (in orange) interval. The red asterisk in the upper left corner of a graph signals a significant time x interval duration interaction. Right side, rescaled curves: the time axis for the 30-s data (in orange) was multiplicatively rescaled. Superposition between the two curves was indexed by eta-square ($\eta^2$) indicated in the bottom right of each graph, the highest values being highlighted in red.*

### Parallel training, Gamma activity

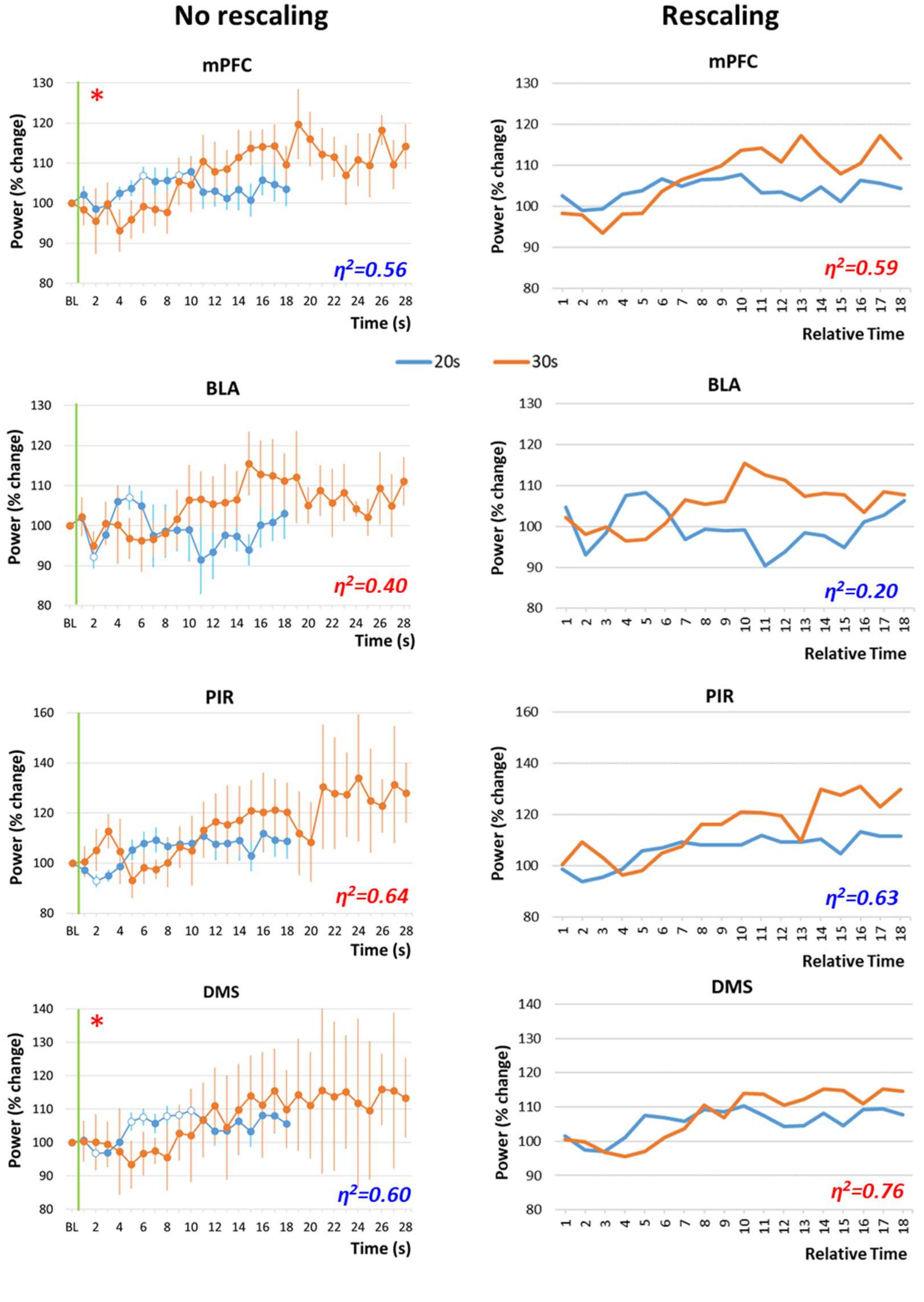


**Supplementary Figure S5:** *Gamma (40-80Hz) band mean power variations in Group 20s and Group 30s, during the odor-shock interval in the four recording sites. mPFC: medial prefrontal cortex; BLA: basolateral amygdala; PIR: olfactory piriform cortex; DMS: dorso-medial striatum. Left side, raw data: gamma power changes are represented from odor onset (green vertical line) during the 20s (in blue) or the 30s (in orange) interval. The red asterisk in the upper left corner of a graph signals a significant time x interval duration interaction. Right side, rescaled curves: the time axis for the 30-s data (in orange) was multiplicatively rescaled. Superposition between the two curves was indexed by eta-square ($\eta^2$) indicated in the bottom right of each graph, the highest values being highlighted in red.*

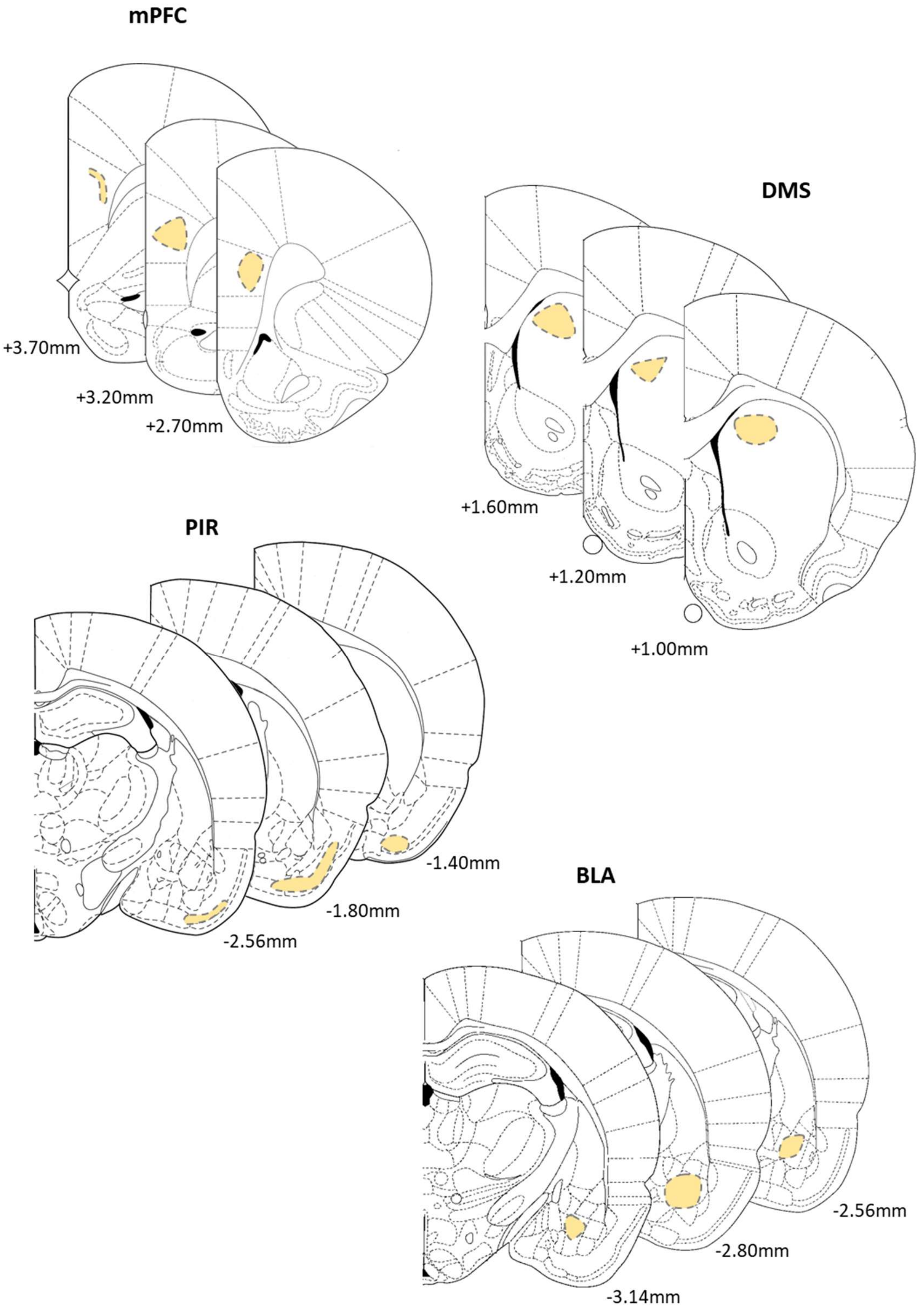


**Supplementary Figure S6:** *Areas targeted by the electrodes (light orange areas) in the three recording sites. Numbers at the bottom indicate the relative position of coronal slices from bregma (Adapted from Paxinos and Watson, 2007). mPFC: medial prefrontal cortex; DMS: dorso-medial striatum; PIR: olfactory piriform cortex; BLA: basolateral amygdala.*
